# Uni-TINT: Unifying T and Innate Lymphoid Cell Taxonomy with a high resolution, pan-disease and pan-tissue single-cell transcriptomic atlas

**DOI:** 10.64898/2026.07.20.739448

**Authors:** Samuel W.J. Chuah, Mengwei Li, Kok Siong Ang, Nicholas R.J. Gascoigne, Jinmiao Chen

## Abstract

T cells and innate lymphoid cells (ILCs) exhibit extensive phenotypic diversity across tissues and diseases, yet inconsistent annotation limits cross-study comparisons and biological interpretation. We present Uni-TINT, an integrated pan-disease and pan-tissue atlas comprising 3.46 million cells from 1,869 samples spanning 194 studies, 166 disease subtypes, and 66 tissue types. Through systematic, hierarchical manual annotation, Uni-TINT establishes a unified and context-aware taxonomy of T and ILC populations, resolving 207 cell subtypes and states across conventional and unconventional T cells, natural killer (NK) cells, helper ILCs, thymocytes, and hematopoietic progenitors. We identified an immunosuppressive, tumour-associated CD4⁺ T regulatory population which we validated with spatial transcriptomics. Other rare and unconventional populations characterised included CD8⁺ regulatory T cells, invariant NKT cells and memory-like NK cells. Integration of T cell receptor sequencing suggested functional associations between γδ T cell co-receptor expression and TRDV gene usage. Finally, a comparative analysis of healthy and diseased immature cells identified a small population of malignant hematopoietic stem cells carrying chromosomal aberrations and enriched in acute leukaemia of mixed phenotype. Together, Uni-TINT provides a unified reference framework for immune annotation and discovery across health and disease.

**Highlights:**

- Uni-TINT harmonises T and ILC nomenclature across diseases and tissues
- Atlas-guided discovery and spatial transcriptomics validation identify a tumour-associated CD4⁺ Treg population
- High-resolution clustering and annotation enable characterisation of rare CD8⁺ Tregs, invariant NKT cells, and memory-like NK cells
- Paired TCR sequencing links γδ T cell states to TRDV gene usage

## Introduction

T cells and innate lymphoid cells (ILCs) are two lymphoid-lineage cell types from adaptive and innate immunity respectively, that play important and overlapping roles. T cell activation is generally antigen-specific^1,2^. The main T cell types include conventional CD4^+^ and CD8^+^ T cells expressing the ⍺β T cell receptor (TCR), and unconventional T cells such as the mucosal-associated invariant T (MAIT) cells, natural killer T (NKT) expressing invariant or semi-invariant TCRs. Additionally, there are *γ*δT cells expressing the alternative *γ*δ TCR. ILCs, on the other hand, rely on antigen-independent activation which is regulated by balance of activating and inhibitory signals delivered by cell surface receptors and cytokines^3^. They encompass both cytotoxic ILCs, also known as natural killer (NK) cells, and helper ILCs^3^. However, the human immune system is extremely complex and diverse, and outside of these clear definitions, T and ILCs have overlapping phenotypes. Some T cells, especially the unconventional subtypes, can display innate-like properties, being activated instead by innate immune signals^4^ and expressing NK receptors which promote cytotoxicity upon activation^1^. Similarly, some NK cell subtypes can display adaptive-like characteristics like clonal-specificity and memory^5^. In addition, researchers have identified helper ILC subsets that share similar transcription factors and cytokine profiles with the CD4^+^ T helper subsets^3^. Both T and ILCs exhibit considerable plasticity in response to the disease and/or tissue context they are in, functioning to protect the host from disease, failing which, these cells can contribute to disease pathogenesis and/or progression. Context-aware annotation of the various cell types and cell states of T and ILCs is thus an essential part of understanding their function and phenotype in health and disease for better clinical decision making and development of new therapies^6^. For the purposes of this study, cell type will be defined as taxonomy that characterises a cell according to its lineage (e.g. CD4^+^, CD8^+^ and NK cells) and sub-lineage or subtype (e.g. Th1, Th2 and Th17 within CD4^+^ T cells), while cell state will refer to taxonomy that describes the differentiated or functional state of the cell (e.g. effector, memory, activated, exhausted, etc.)^7,8^. Thus, a cell type can exist in different cell states, depending on the local tissue context and the broader disease context that the cell is in.

In the past, immune profiling with low-dimensional technologies was limited to broad categories of subtypes, relying on a few protein markers or secretion profiles to characterise cells^9^. The development of high-dimensional technologies such as single cell RNA sequencing (scRNAseq) has made immune profiling both richer and more complicated^6,9^. With scRNAseq, nuances in phenotype and transitional states between cells can be captured, which is highly valuable in characterising rare cell types and states involved in a disease or condition^6^. Indeed, with scRNAseq becoming increasingly affordable and accessible, individual immune profiling studies can now achieve extremely in-depth analyses of immune subtypes of interest. They engage in context-aware phenotyping and try to understand the function of a particular immune cell type in light of the disease and/or tissue environment where it was found^6^. However, individual studies are often limited in sample size and cell numbers, reducing the resolution of analysis less comprehensive and potentially obscuring the identification of rare or transitional immune subtypes. They also tend to specialise in a particular disease or condition, making any application beyond the studied condition challenging.

Large-scale atlases help alleviate such limitations by collecting and compiling large numbers of studies into a single resource, providing access to samples and conditions otherwise not available^10^. Various specialised immune cell atlases have been built, focusing on different conditions like healthy subjects^11^, immunotherapy in cancer^12,13^, and SARS-COV-2^14^. Others focus on a particular immune cell type like T cells^15–17^, B cells^18,19^, NK cells^20^, and myeloid cells^21–23^, as well as the adaptive immune repertoires – the T cell receptor (TCR) and B cell receptor (BCR)^15,24–26^. Nevertheless, while these represent a large improvement over individual studies, current immune atlases still tend to be restricted in either disease or tissue context. Those which contain samples from multiple tissues or organs often focus on a particular disease or on healthy samples, while those offering a larger number of diseases tend to be restricted in terms of tissue of origin, for example, the blood. These also tend to specialise in a particular category of disease, for example cancer or autoimmune diseases. Furthermore, the cell type annotations that current immune atlases offer tend to be less granular than those available in an individual study, with broader classifications that are less context-aware. Finally, a common limitation in the immune profiling of T and ILCs, both in individual studies and in atlases, is the lack of consensus in the definitions and annotation of cell type and state^9^. For example, atlases commonly annotate CD4^+^ T cells by subtypes (e.g. Th1, Th2, and Th17) while CD8^+^ T cells are commonly annotated by cell state^1^, even though many individual studies have described CD8^+^ T cell subtypes which are analogous to at least some of those in the CD4^+^ population^27,28^. Considerable flexibility and inconsistencies also exist, partially due to the inherent plasticity of T and ILCs, i.e. the fact that each cell subtype studied takes on a different phenotype depending on its disease and tissue context. This makes immune cell analyses and comparisons inconvenient and largely restricted to similar diseases or tissues of origin. There is thus a need for immune cell atlases with harmonised and hierarchical annotations of cell types and cell states, with highly granular clustering and phenotyping at a resolution comparable to individual studies, and with comprehensive disease and tissue coverage and integration for broad applicability and analyses.

To address these needs, we constructed Uni-TINT (**Uni**fying **T** and **IN**nate lymphoid cell **T**axonomy), a pan-disease and cross-tissue single-cell atlas of T and ILCs, aimed at refining and harmonising T and ILC annotations, using publicly available datasets compiled in DISCO^29,30^, along with manually curated metadata of sample-related information including disease, tissue, age, and Human Leukocyte Antigen (HLA) haplotype. We conducted fine clustering and manual annotation of immune cell populations, following a systematic hierarchy of the main immune cell lineages, followed by increasingly finer subtypes and cell states within the subtypes, yielding most of the expected subtypes and cell states. We also incorporated disease and tissue context in our annotations to enhance the understanding of cell type and function, using *in silico* tools and methods as a first level of validation where possible. With our fine-grained annotations, we described new subtypes of CD4^+^ regulatory T cells (Tregs) in new disease conditions, identified rare cell types like CD8^+^ Tregs and invariant NKT (iNKT) cells, characterised unconventional and experimental cell types such as memory-like NK cells in primary human samples, and used cross-study and cross-disease analyses to identify a malignant hematopoietic stem cell (HSC) cluster. The data used to construct Uni-TINT also contained samples with associated single-cell TCR sequencing (scTCRseq) data, which we used to explore associations between gene expression and TCR usage, providing insights into a potential functional link between co-receptor expression and TRDV gene usage in *γ*δ T cells. Finally, we employed spatial transcriptomics data to show that our atlas’s fine grained and context-aware annotations using scRNAseq successfully identified a solid cancer-associated CD4^+^ Treg cluster, whose tissue-infiltrating and immunosuppressive properties could be validated in tumour tissue samples.

## Results

### Construction of a pan-disease, pan-tissue single-cell transcriptomic atlas of T and ILCs

We first accessed DISCO to obtain all publicly available 10x Genomics scRNAseq data with associated metadata. We removed samples sorted for non-T and ILCs, as well as genetically modified samples, retaining 1869 samples belonging to 194 projects [Table S1]. These samples encompass 166 disease subtypes (14 disease groups) and 66 tissue types (12 tissue groups) [Figure 1A, Table S2 and S3]. Where available, matching scTCRseq data was also downloaded [Table S1]. After stringent sample selection and quality control, we extracted the T and ILCs and performed atlas level integration, incorporating the associated scTCRseq data [Figure 1B, Methods]. The integrated data was then annotated in a multi-step process.

**Figure 1.**
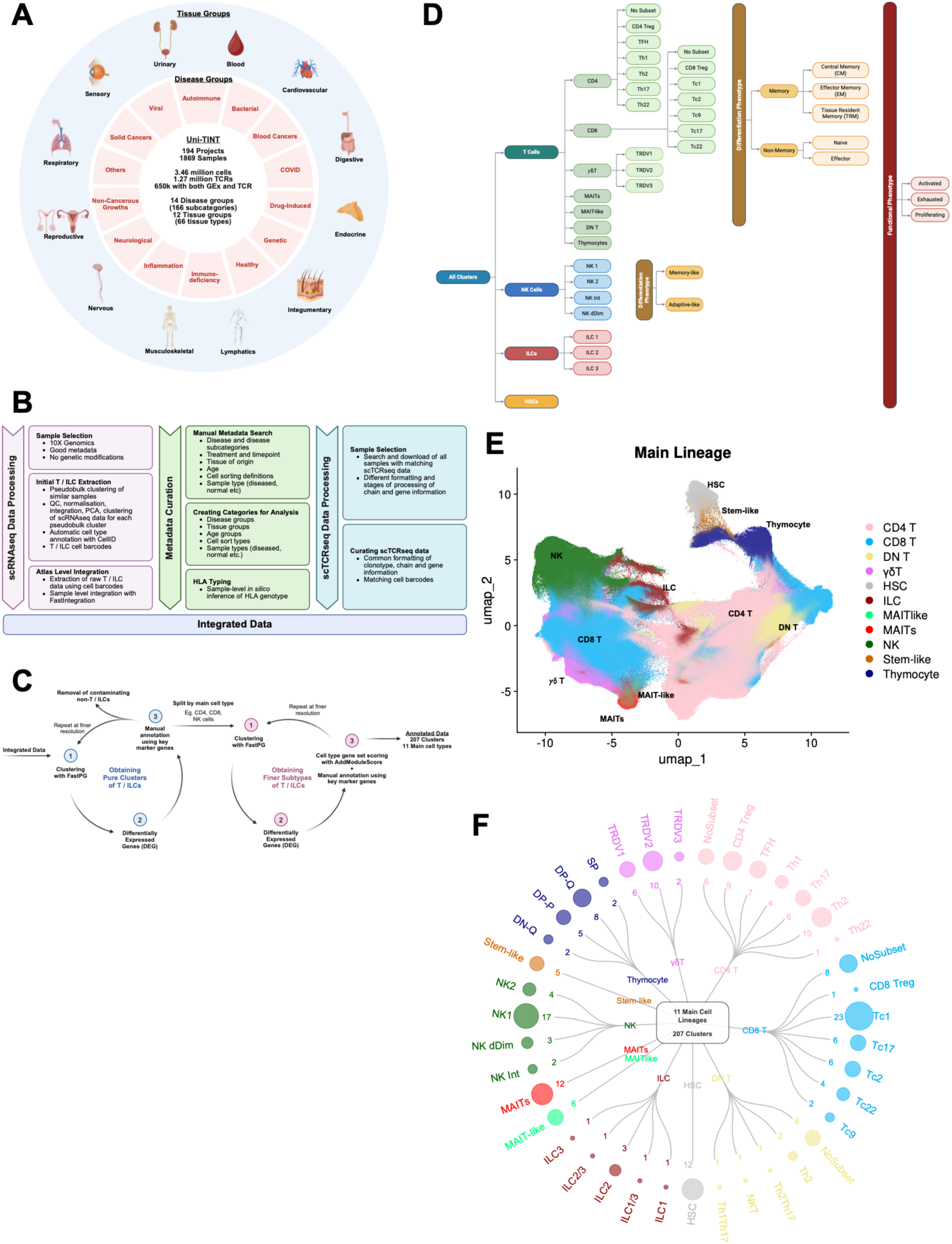
Construction of a pan-disease, pan-tissue single-cell transcriptomic atlas of T and ILCs. (**A**) Schematic summarising the atlas and its disease and tissue groupings. (**B**) Flowchart detailing the process of constructing the atlas. (**C**) Schematic describing the two-staged manual annotation process. (**D**) Flowchart showing the systematic, hierarchical manual annotation process. (**E)** Uniform manifold approximation and projection (UMAP) visualisation all clusters, coloured by main cell lineage. (**F**) Circular dendrogram showing main cell lineages and the subtypes within each lineage. The numbers and size of the circles indicate the number of individual clusters within each subtype. Figure parts A-C created with BioRender.com.

We adopted a two-staged, reiterative process to achieve high resolution separation and labelling [Figure 1C, Methods]. The first stage aimed to achieve pure clusters of T and ILCs. The cells were clustered and computed for differentially expressed genes (DEGs) and manually annotated based on marker genes [Table S4]. Where found, we removed any non-T cell and non-ILC clusters. This process was repeated at finer resolutions until only pure clusters of T and ILCs remained [Figure 1C]. The second step aimed to generate granular clusters of the various subtypes of T and ILCs. We first split the data into the major cell types, namely T cells, NK cells, ILCs, and others. We then iteratively clustered the cells, computed the DEGs and gene module scores, and manually annotated each cluster based on marker genes [Table S4]. The annotation encompassed main cell lineage, cell subtype, memory status, differentiation phenotype, and functional phenotype [Figure 1D]. At the highest level of main lineage, there were 11 main lineages of conventional CD4^+^ and CD8^+^ ⍺β T cells, unconventional T cell lineage MAIT/MAIT-like cells, *γ*δ T cells, and double-negative (DN) T cells, NK cells, helper ILCs, immature thymocytes and hematopoietic stem cells (HSCs) [Figure 1E, F]. At the finest clustering level, there were 207 clusters with assigned subtypes and memory, differentiation and functional phenotypes. Details of the manual annotation process are provided in the Methods section. In its final form, the T/ILCs atlas has 3.46 million cells in 207 clusters.

To facilitate context-aware phenotyping in the downstream analyses, we manually curated and harmonised the metadata from each sample and project, condensing available patient demographic information, including disease, treatment, tissue of origin, sorting strategy and age, into categories for analysis [Figure 1B, Table S1, Methods]. Inference of HLA allele was also carried out *in silico* using arcasHLA^31^ and visualised according to HLA class and frequency in the various disease groups [Figure S1C, D].

### Disease- and tissue-focused characterisation of CD4^+^ T cells reveals new subsets in new contexts

As the “helper” compartment of adaptive immunity, CD4^+^ T cells play important roles in regulating and coordinating immune responses, especially the functions of other immune cells^1^. The CD4^+^ T cell subtypes are generally well-defined in terms of their transcription factor programs and secretory profiles^1^. Therefore, we focused on CD4^+^ T cells first to show that our fine clustering and manual annotation process could produce the known CD4^+^ T cell subtypes. At its finest level of clustering, our atlas has 43 CD4^+^ T cell clusters. Depending on the confidence of annotation based on canonical markers found in the DEGs and gene module scoring, we assigned either a primary or secondary subtype (or both) [Table S4, Methods], with six subtype labels in total, namely T regulatory (Treg), T follicular helper (TFH), and T helper (Th) subtypes 1, 2, 17, and 22 [Figure 2A, B]. A small number of clusters did not fall into any existing classification and were assigned to NoSubset. These were generally of naïve or transitioning phenotypes. We were unsuccessful at identifying Th9 cells in our data. For all clusters, we also assigned a memory status, differentiation phenotype, and functional phenotype [Figure 2A, C, S7A, Methods]. To provide disease and tissue context to each cluster, disease and tissue group proportions were visualised by cluster [Figure 2A, S7B-E] and we tested for enrichment in each group using Fisher’s exact test, showing only the significantly enriched clusters [Figure 2D, E]. To demonstrate the usefulness of our annotations to facilitate disease and tissue context-aware phenotyping, we selected the well-studied CD4^+^ Tregs for in-depth characterisation to see if biologically meaningful insights could be drawn for each CD4^+^ Treg cluster, congruent with its disease and tissue of origin.

**Figure 2.**
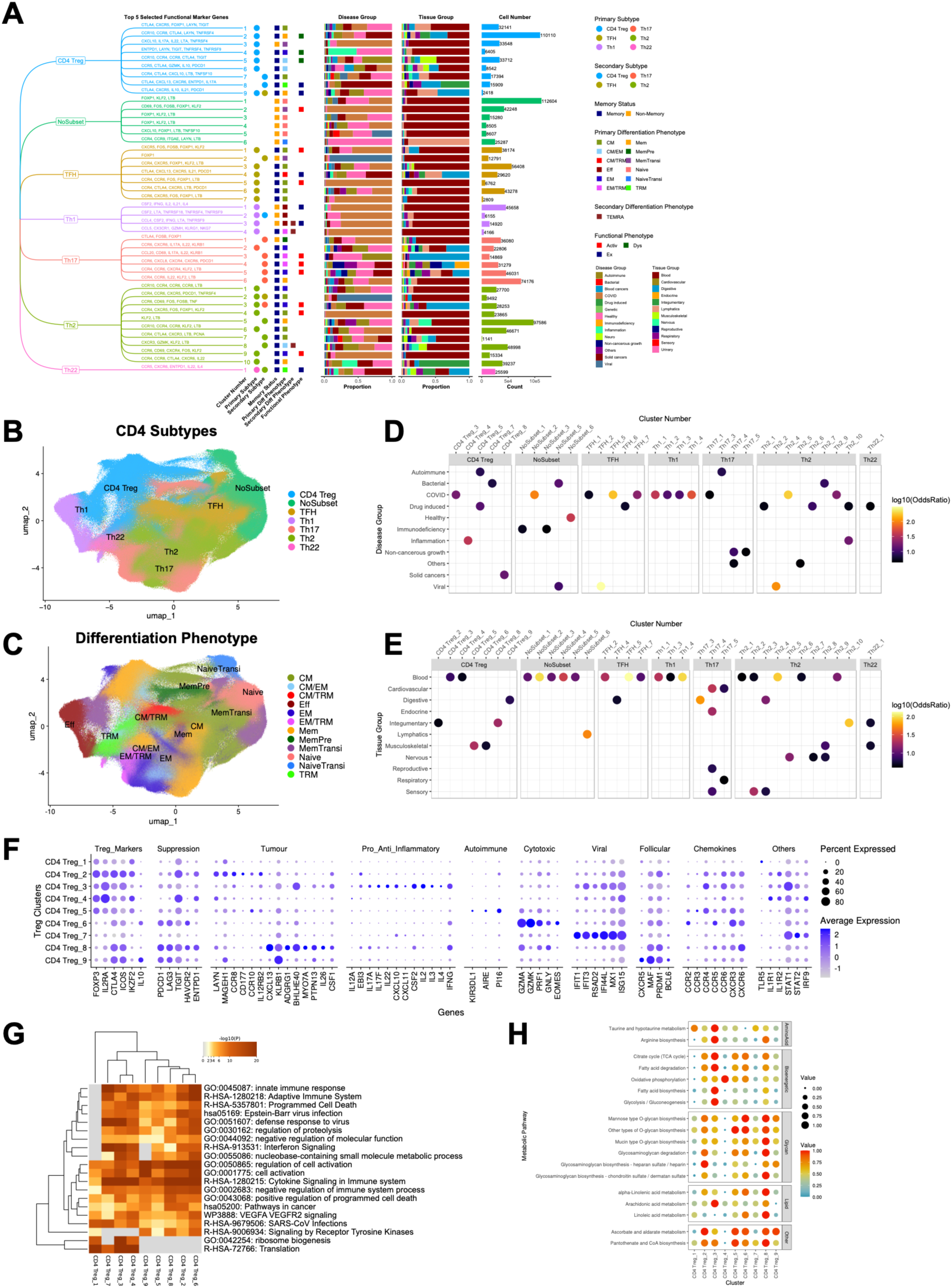
CD4^+^ T cell compartment. (**A**) Tree diagram showing the CD4 subtypes, clusters belonging to each subtype (named by number) and the top five selected functional marker genes of each cluster. The primary and/or secondary subtype annotations of each cluster are indicated by the coloured circles, while the memory status, primary and/or secondary differentiation phenotype and functional phenotypes are indicated by coloured squares. Stacked bar plots show the proportion of cells of each cluster coming from the various disease and tissue groups, with the cell numbers plotted in bar plots beside, coloured by subtype. (**B**) UMAP of CD4 subtypes, coloured by subtype. (**C**) UMAP of CD4 subtypes, coloured by differentiation phenotype. (**D**) and (**E**) Dot plot heatmaps showing CD4 clusters significantly enriched in the various disease and tissue groups respectively [P < 0.05 after Bonferroni correction and Odds Ratio (OR) > 4]. (**F**) Dot plot showing expression levels of selected genes for CD4 Treg clusters. (**G**) Top 20 enriched functional pathways across CD4 Treg clusters generated by Metascape, based on differentially-expressed genes (DEGs) calculated against all cells. (**H**) Selected metabolic pathways enriched across CD4 Treg clusters, computed using scMetabolism.

There were nine clusters of CD4^+^ Treg cells with diverse phenotypes, of which three clusters were found to contain cells from a heterogeneous mix of disease groups, without being significantly enriched in any [Figure 2A (Disease Group barplots), 2D]. The first cluster, CD4_Treg_1, potentially regulates inflammation of bacterial origin with its expression of *LAYN* and *MAGEH1* [Figure 2F] which describe inflammation-exposed Tregs^32^, and *TLR5* which enhances the suppressive capacity of Tregs when exposed to bacterial flagellin^33^. The second, CD4_Treg_6, highly expressed *EOMES*, *IL10* and *PDCD1*, while having lower *FOXP3* expression than the other Treg clusters. It also had a higher expression of cytolytic genes *GZMA*, *GZMK*, *PRF1* and *GNLY*. It is therefore likely to be a cytotoxic Type 1 regulatory (Tr1) subtype which exerts its immunosuppressive function in various ways, including killing antigen-presenting cells or other T cells^34^. The third, CD4_Treg_9, showed characteristics of a T follicular regulatory (Tfr) subtype with its expression of *CXCR5* together with canonical Treg genes, but lower *FOXP3* expression^35^, along with the genes *BCL6*, *PRDM1* and *MAF*, which are important for Tfr development^36^. Finally, CD4_Treg_7, while significantly enriched in Bacterial diseases, also comprised a heterogenous mix of disease groups. It expressed an “acute respiratory viral signature” (*IFIT1*, *IFIT3*, *RSAD2*, *IFI44L*, *MX1*, *ISG15*) [Figure 2F] which has been associated with rhinovirus infection of cells^37^, as well as *STAT1*, *STAT2* and *IRF9* [Figure 2F], which together indicate activation by type I IFN stimulation, a process that can be triggered by viral infections^38^.

Of the remaining clusters, three had cells derived from specific diseases or tissue sources. The first was CD4_Treg_3 that were mostly derived from COVID blood samples [Figure 2A, S7D]. This cluster appeared to be a potentially dysregulated and inflammatory Treg subtype that expressed both pro-inflammatory and anti-inflammatory cytokines [Figure 2F], potentially contributing to COVID pathology [Figure 2D, E, S7D, E]. It expressed the inflammatory cytokines, *IL17A, IL17F, IL3, CSF2, CXCL10*, and *CXCL11*^39–41^, of which *CXCL10* and *CXCL11* have been observed to be elevated in the serum of COVID patients^42^. It also expressed the anti-inflammatory cytokines *IL4*^43^ and IL35, which is formed from the gene products of *IL12A* and *EBI3*^44^ [Figure 2F]. Finally, it expressed *IL22*, suggesting *FOXP3* dysregulation because *IL22* is a direct target gene repressed by *FOXP3*^45^. Interestingly, it also expressed the acute respiratory viral signature and type I IFN stimulation signature described in CD4_Treg_7 above, suggesting that both clusters might have arisen during a respiratory virus infection. The second cluster, CD4_Treg_4, was primarily derived from blood samples of healthy subjects, and subjects with inflammatory diseases [Figure 2A, D, E], implying a role in regulating inflammatory conditions. It expressed *IL1R1* and *IL1R2*, a characteristic of activated Tregs^46^, *LAYN* and *CCR8* which suggests exposure to an inflammatory environment^32^, and *TIGIT* which plays a role in inhibiting Th1 and Th17 inflammatory responses^47^ [Figure 2F]. The third cluster, CD4_Treg_5, was significantly enriched in autoimmune diseases and drug-induced conditions, with a large proportion of its cells originating from autoimmune diseases [Figure 2A, D, E]. It expressed *KIR3DL1* and *PI16* which inhibit Treg function and/or differentiation^48,49^, suggesting a pathogenic role of reduced Treg function in autoimmunity. It also expressed *AIRE* [Figure 2F], which has been described to direct autoreactive T cell types into the FOXP3 lineage^50^, suggesting a possible origin for these cells.

The last two clusters, CD4_Treg_2 and CD4_Treg_8, were associated with solid cancers, with statistical significance for CD4_Treg_8 [Figure 2A, D]. CD4_Treg_2 expressed markers which have been associated with tumour infiltrating Tregs, namely *CD177*^51^ and *LAYN*, *MAGEH1*, and *CCR8*^32^ [Figure 2F], along with chemokine receptors such as *CCR10* and *CXCR6*^52,53^ [Figure 2F]. CD4_Treg_8 on the other hand expressed a gene signature (*CXCL13*, *KLRB1*, *ADGRG1*, *BHLHE40*, *MYO7A*, and *PTPN13*) [Figure 2F] which has previously been described in activated, highly differentiated, and suppressive Tregs found in the synovial fluid of patients with juvenile idiopathic arthritis^54^. It also expressed higher levels of *PDCD1, LAG3, TIGIT and HAVCR2*, all of which contribute to the immunosuppressive properties of FOXP3^+^ CD4^+^ Tregs^55^. Finally, it expressed *CSF1*, a ligand for tumour-associated macrophages and associated with poorer cancer prognosis^56^. Thus, CD4_Treg_8 represents a known Treg phenotype that may be implicated in a new disease context. Consequently, we investigated the CD4_Treg_8 cluster further.

We first examined CD4_Treg_8’s association with diseases, especially cancer. Interestingly, the cluster comprised of cells from at least six main kinds of solid cancers, breast cancer (BRCA), basal cell carcinoma (BCC), hepatocellular carcinoma (HCC), non-small cell lung cancer (NSCLC), nasopharyngeal carcinoma (NPC), and squamous cell carcinoma (SCC) [Figure S7D]. One commonality among these cancers is that they are all carcinomas involving epithelial cell or epithelial cell-like malignancy. Notably, the two other primary diseases which contributed to the CD4_Treg_8 cluster were COVID and ulcerative colitis, which also involve epithelial cell pathology. To characterise CD4_Treg_8 further, we performed over-representation analysis (ORA) using Metascape^57^. CD4_Treg_8 showed enrichment in pathways related to cancer, regulation of cell activation, and negative regulation of immune system processes, highlighting it as one of the most immunosuppressive among the Treg clusters [Figure 2G]. Finally, we examined the metabolic profile of CD4_Treg_8 with scMetabolism^58^ and found that it showed the highest enrichment of O-glycan biosynthesis among the Treg clusters [Figure 2H], which plays a role in controlling T cell trafficking^59^. It also had high enrichment of glycosaminoglycan biosynthesis of chondroitin/dermatan sulphate, which is controlled by the enzyme dermatan sulphate epimerase (DSE). While DSE expression has not been described in Tregs, it has been linked to enhanced anti-tumour immunity and infiltration of CD8^+^ and CD4^+^ T cells^60^, and could potentially enhance the infiltration of Tregs too. Taken together, these results show that CD4_Treg_8 is a tissue-infiltrating Treg cluster recruited to diseased or inflamed epithelial tissue where it potentially exerts immunosuppressive functions. It is therefore a potential treatment target for reducing immunosuppression in solid tumours, especially those of epithelial origin.

### Fine-grained characterisation of CD8^+^ T cells reveals rare populations

CD8^+^ T cells form another compartment of adaptive immunity and are generally responsible for eradicating pathogen-infected cells and abnormal cells such as malignant tumour cells^1^. Compared to CD4^+^ T cells which are more clearly defined by their subtypes, the annotation of CD8^+^ T cells is more varied, and they are more commonly described in terms of their differentiation state, such as effector versus memory^1^. However, studies have shown that CD8^+^ T cells can also be categorised into subsets based on their transcription factor and secretory profiles, analogous to the CD4^+^ Th subtypes^27^.

To show that our fine clustering and manual annotation process could also incorporate the diverse definitions of CD8^+^ T cell subtypes and cell states, we annotated the CD8^+^ T cell clusters in our atlas in a similar way to CD4^+^ T cells. The atlas has 51 CD8^+^ T cell clusters which are annotated with the subtypes of Tc1, Tc2, Tc9, Tc17, Tc22, CD8^+^ Tregs and NoSubset [Figure 3A, B], and labelled with differentiation, memory, and functional phenotypes [Figure 3A, C, S8A], based on manual examination of DEG lists for gene markers and gene module scoring [Table S4, Methods]. Disease and tissue group proportions were visualised by cluster [Figure 3A, S8B-E] and cluster enrichment in each group was computed using Fisher’s exact test [Figure 3D and E] to provide disease and tissue context.

**Figure 3.**
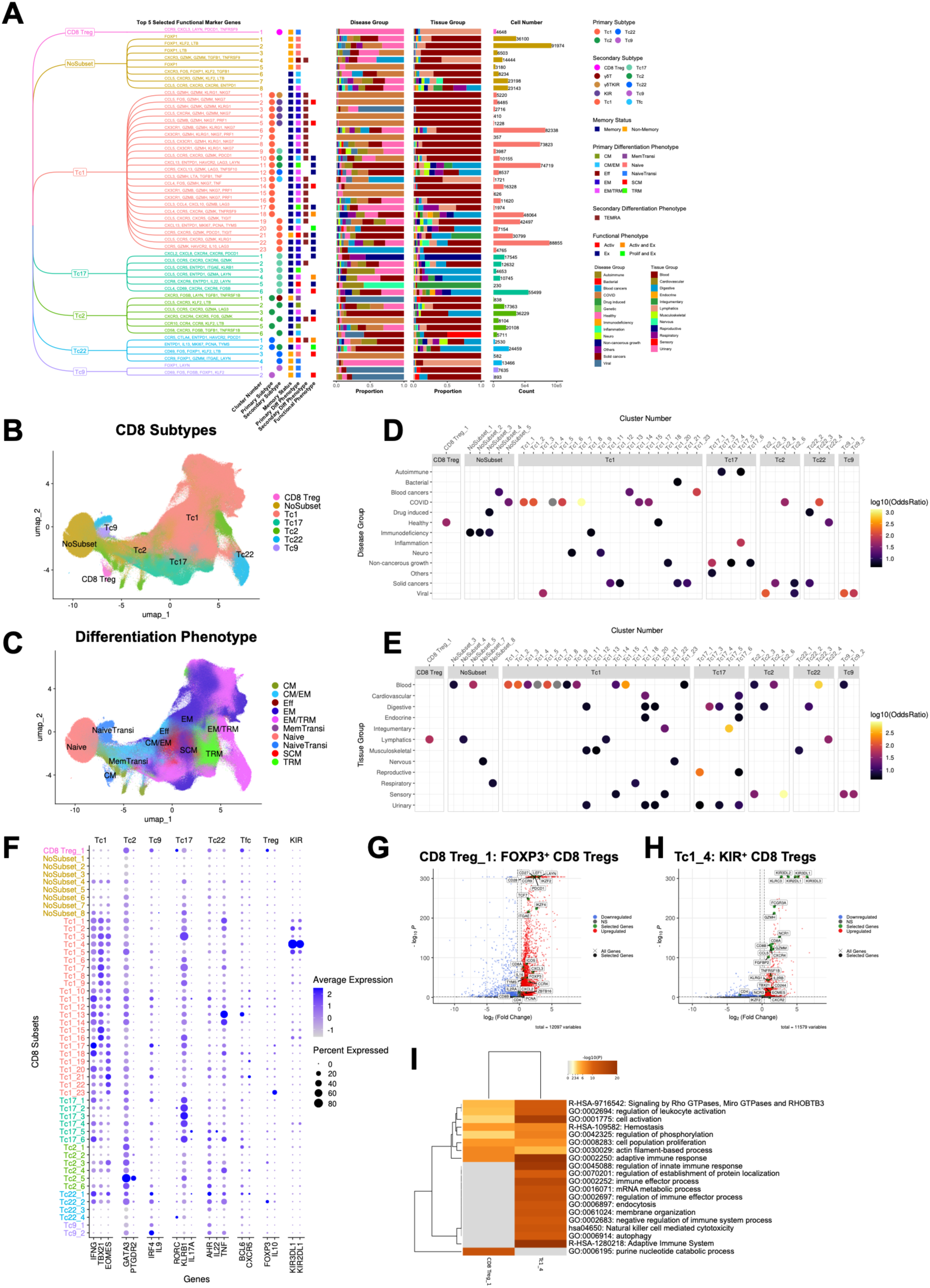
CD8^+^ T cell compartment. (**A**) Tree diagram showing the CD8 subtypes, clusters belonging to each subtype (named by number) and the top five selected functional marker genes of each cluster. The primary and/or secondary subtype annotations of each cluster are indicated by the coloured circles, while the memory status, primary and/or secondary differentiation phenotype and functional phenotypes are indicated by coloured squares. Stacked bar plots show the proportion of cells of each cluster coming from the various disease and tissue groups, with the cell numbers plotted in bar plots beside, coloured by subtype. (**B**) UMAP of CD8 subtypes, coloured by subtype. (**C**) UMAP of CD8 subtypes, coloured by differentiation phenotype. (**D**) and (**E**) Dot plot heatmaps showing CD8 clusters significantly enriched in the various disease and tissue groups respectively [P < 0.05 after Bonferroni correction and Odds Ratio (OR) > 4]. (**F**) Dot plot showing expression levels of selected genes for all CD8 clusters, grouped and coloured by subtype. (**G**) and (**H**) Volcano plots showing differentially-expressed genes (DEGs) of the two CD8 Treg clusters (CD8 Treg_1 and Tc1_4 respectively), calculated against all other cells. (**I**) Top 20 enriched functional pathways across CD8 Treg clusters generated by Metascape, based on differentially-expressed genes (DEGs) calculated against all cells.

Compared to the CD4^+^ subtypes, the CD8^+^ subtypes were more ambiguous and harder to annotate. For example, the CD8^+^ clusters would either express markers and transcription factors of more than one subtype, or did not express the full set of marker genes of a particular subtype [Figure 3F, S5B]. Therefore, the former case was resolved by assigning a primary and secondary subtype while the latter case would only have a secondary subtype [Table S4, Methods]. This subtype ambiguity demonstrates the plasticity of CD8^+^ T cells and could indicate their ability to adapt to their local tissue environment^27^.

In the process of annotating the CD8^+^ clusters, we discovered two clusters which fit the definitions of CD8^+^ Tregs. These are less well-studied compared to CD4^+^ Tregs and are heterogeneously defined, being primarily identified by an immune regulatory function rather than a specific set of marker genes^61,62^. Due to their rarity, they are difficult to isolate in studies with limited sample sizes. FOXP3^+^ CD8^+^ Tregs, for example, comprise merely 0.1 to 1% of total CD8^+^ T cells in the blood^62^. However, the large sample and cell numbers in our atlas enabled us to identify and characterise them.

The first cluster, CD8_Treg_1 (4648 cells), contained *FOXP3*^+^ CD8^+^ Treg cells originating primarily from healthy thymic tissue [Figure 3A, D, E, S8D, E]. It expressed canonical Treg markers such as *FOXP3*, *IKZF2*, *PDCD1*, and *ICOS* [Figure 3G], and *IL2RA* (only relative to other CD8^+^ clusters) [Figure S8F, Table S6]. It also expressed transcription factors of other CD8^+^ subtypes [Figure 3F] but they were not significant DEGs [Table S5] and hence were not considered for annotation. Another notable expressed gene was *ITGAE* (CD103), which supported its thymic origin^61^. It also displayed some markers of naïve T cells (*TCF7*, *LEF1*, *CD27*, and *CD28*, but without *SELL* and *CCR7*), together with *CCR9* [Figure 3G, S8F], a receptor expressed by both immature and mature thymocytes^63^, and *TYMS* and *PCNA*, markers of proliferation. Therefore, we postulate that this was a cluster of mature naïve T cells transitioning out of the thymus. Finally, the *CD8A* gene was more highly expressed than *CD8B* [Figure 3G, S3A], and *ZBTB16* (PLZF) was more highly expressed relative to the other CD8^+^ clusters [Figure S8F, Table S6], suggesting that these could also be CD8⍺⍺ Tregs^62^. To our knowledge, FOXP3^+^ CD103^+^ CD8^+^ Tregs from the thymus have been previously described using flow cytometry^64^, but not at the transcriptomic level, even in human thymic atlases or studies^65,66^. Hence, we believe this is the first successful identification of human *FOXP3*^+^ CD8^+^ Tregs from the healthy thymus at the single cell transcriptomic level.

The other CD8^+^ Treg cluster (410 cells) consisted of Killer cell Immunoglobulin-like Receptor genes positive (*KIR^+^*) CD8^+^ Treg (Tc1_4) cells that were exclusively derived from COVID blood samples [Figure 3A, D, E]. These did not express *FOXP3* but expressed other Treg-associated genes such as *TNFRSF1B* and *IKZF2* at levels higher than other CD8^+^ clusters [Figure 3H, S8G, Table S6]. They also displayed a Tc1 phenotype with the expression of *TBX21* (T-bet) and *EOMES*, high expression of *KIR* genes, effector genes like *FCGR3A* (CD16) and *GZMH*, and NK-associated genes such as *KLRC3*, *NCR1*, and *NCR3* [Figure 3F, H, S8G]. A similar transcriptional profile has been previously described in autoimmune diseases and COVID where such *KIR*^+^ CD8^+^ Tregs were shown to play a role in suppressing self-reactive, pathogenic CD4^+^ T cells through direct killing^67,68^.

We also characterised the two clusters with functional pathway enrichment analysis using Metascape [Figure 3I]. The *FOXP3*^+^ CD8^+^ Treg cluster displayed less activation and slight proliferation, consistent with its transitioning naïve phenotype. On the other hand, the *KIR*^+^ CD8^+^ Treg cluster showed more activation and cytotoxicity, consistent with its function in suppressing other T cells [Figure 3I]. Thus, the fine clustering and detailed annotation of our atlas has enabled us to identify and characterise two rare CD8^+^ Treg populations with disease and tissue context awareness.

### Combined gene expression and TCR analyses reveal functional associations between TCR gene usage and co-receptor expression in *γ*δT cells

Unlike the conventional CD4^+^ and CD8^+^ T cells, unconventional T cells do not respond to peptide antigens presented on the classical Major Histocompatibility Complex (MHC) molecules^69^. Instead, they recognise a diverse range of other antigens like lipids and metabolites that are presented on non-polymorphic antigen-presenting molecules^69^. These T cells tend to have a more restricted TCR and often display innate-like capabilities for rapid immune responses^69^. In this atlas, we annotated four main lineages of unconventional T cells, namely, MAIT, MAIT-like, *γ*δ T, and double negative (DN) T cells [Figure 4A, B]. The DN T cells were labelled as such due to the lack of gene expression of both CD4 and CD8 co-receptors. MAIT and MAIT-like cells were further categorised by *CD4* and *CD8A/B* co-receptor expression, while the *γ*δ T cell clusters were first subdivided by *TRDV1*, *TRDV2*, and *TRDV3* expression, followed by *CD4* and *CD8A/B* expression. As before, all unconventional T cell clusters were annotated with differentiation, memory, and functional phenotypes [Figure 4A, D, S9A] based on their expression of key marker genes [Table S4, Methods]. We also visualised the disease and tissue group proportions per cluster [Figure 4A, S9B-E] and computed cluster enrichment in each group using Fisher’s exact test [Figure 4E and F].

**Figure 4.**
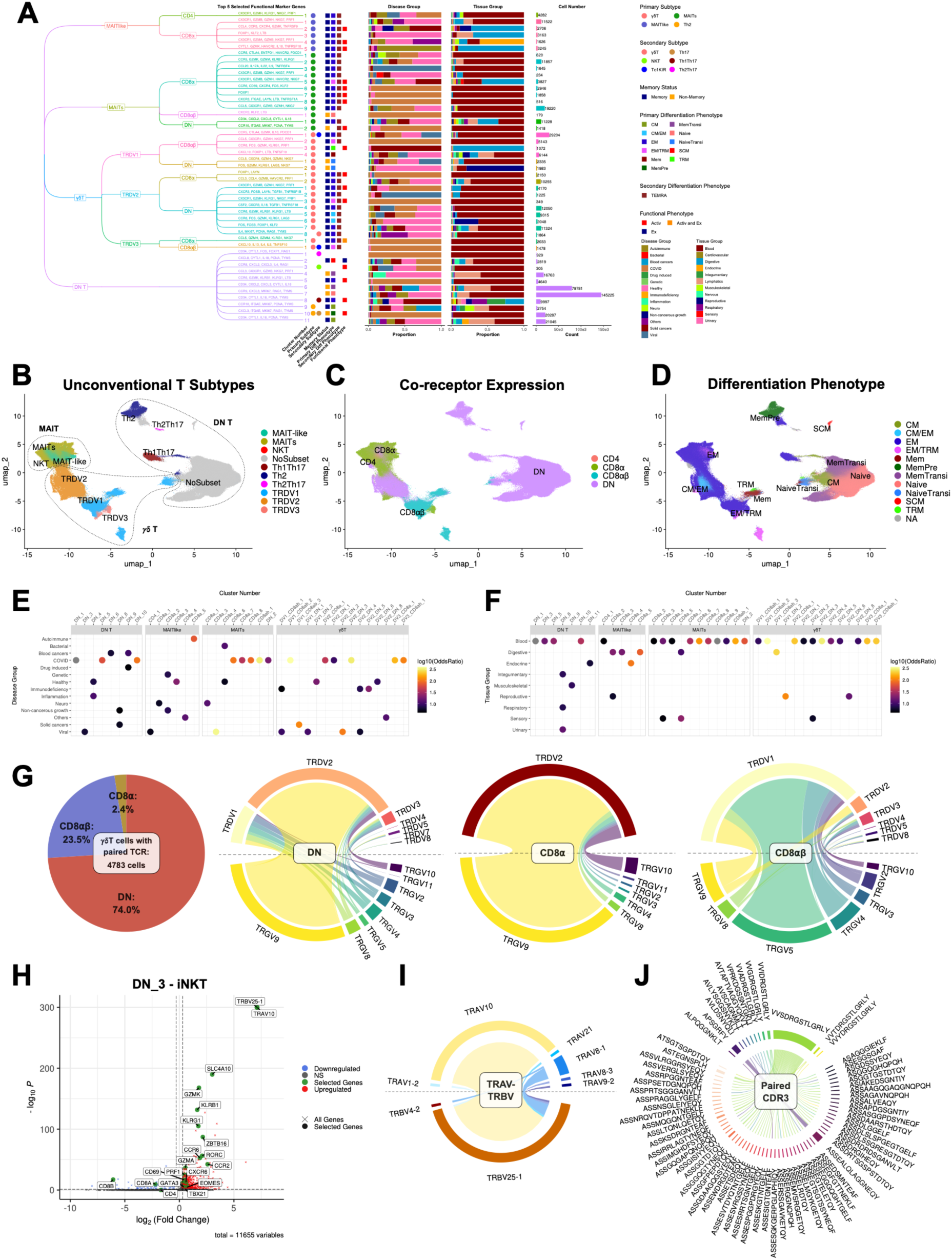
Unconventional T cell compartment. (**A**) Tree diagram showing the Unconventional T cell lineages and their subtypes, the clusters belonging to each subtype (named by number) and the top five selected functional marker genes of each cluster. The primary and/or secondary subtype annotations of each cluster are indicated by the coloured circles, while the memory status, primary and/or secondary differentiation phenotype and functional phenotypes are indicated by coloured squares. Stacked bar plots show the proportion of cells of each cluster coming from the various disease and tissue groups, with the cell numbers plotted in bar plots beside, coloured by subtype. (**B**) UMAP of Unconventional T cell lineages, split by main lineage, coloured by subtype. (**C**) UMAP of Unconventional T cell lineages, coloured by co-receptor expression. (**D**) UMAP of Unconventional T cell lineages, coloured by differentiation phenotype. (**E**) and (**F**) Dot plot heatmaps showing Unconventional T cell clusters significantly enriched in the various disease and tissue groups respectively [P < 0.05 after Bonferroni correction and Odds Ratio (OR) > 4]. (**G**) Circos plots of annotated *γ*δT cells with paired TCR data showing paired TRDV-TRGV gene expression and split by co-receptor expression. (**H**) Volcano plot showing DEGs of the iNKT cluster (DN_3), calculated against all other cells. (**I**) and (**J**) Circos plots showing paired TRAV-TRBV gene expression and paired CDR3⍺-CDR3β respectively in the iNKT cluster (DN_3).

Human MAIT cells express a semi-invariant TCR consisting of *TRAV1-2* rearranged to *TRAJ33*, paired with beta chain genes from the *TRBV6* family and *TRBV20-1*^70^. They typically recognise bacterial metabolite antigen presented on the MHC-related protein 1 (MR1)^70^. In this atlas, we annotated 18 clusters of MAIT and MAIT-like cells [Figure 4A]. The MAIT-like label was assigned to cells that lacked gene expression of the semi-invariant TCR, but otherwise highly expressed the MAIT cell markers. MAIT and MAIT-like clusters generally had similar gene expression profiles [Figure S9F] and clustered together on the UMAP plot [Figure 4B]. They expressed high levels of *ZBTB16* (PLZF) and *KLRB1* (CD161), which are markers of innate immunity^71,72^, and other effector genes such as *PRF1*, *IFNG*, *TNF*, *GNLY* and *KLRG1*, with variation in expression between clusters rather than between subtypes [Figure S9F]. They also generally did not express appreciable levels of the *KIR* genes [Figure S9F]. MAIT and MAIT-like cells were then further classified by their expression of *CD4*, *CD8A* and *CD8B* [Figure 4A, C, S3A]. As expected, most of the clusters expressed either *CD8A* alone, or *CD8A* and *CD8B* together [Figure 4A, C]^70^. Two clusters were negative for both *CD4* and *CD8A/B* while one cluster was *CD4* positive. The CD4^+^ MAIT-like cluster was annotated as such because while it expressed the MAIT cell markers and *TRAV1-2*, the cluster also co-expressed a variety of other TCR genes and thus did not appear to consist purely of CD4^+^ MAITs [Figure S9J].

The next class of unconventional T cells in this atlas were the *γ*δT cells, named after the alternative gamma and delta TCR chains that they express. They are generally considered to be non-MHC restricted because their antigen recognition does not depend on processing by antigen-presenting cells and subsequent presentation on MHC molecules^73^. Human *γ*δT cells are typically classified into three major subtypes based on the *TRDV* gene they express, namely *TRDV1*, *TRDV2*, or *TRDV3*. The *TRDV2* gene is often paired with *TRGV9* and this forms the largest group of *γ*δT cells, followed by *TRDV1* and *TRDV3*, both of which have more unrestricted *TRGV* gene pairing^73^. In this atlas, 18 clusters of *γ*δT cells were identified [Figure 4A, B]. Classifying them based on the gene expression of their *TRDV* and *TRGV* genes, there were 10 TRDV2, 6 TRDV1, and 2 TRDV3 clusters [Figure 4A]. Consistent with the general classification, the TRDV2 clusters also predominantly expressed *TRGV9*, while the TRDV1 and TRDV3 clusters were more heterogeneous in their *TRGV* expression [Figure S9G]. Moreover, the TRDV2/GV9 clusters showed more innate-like expression profiles with higher expression of *ZBTB16*, *KLRB1*, and *IL18RAP*^71,72,74^ [Figure S9G]. On the other hand, the TRDV1 and TRDV3 clusters showed higher levels of *ZNF683*, a marker associated with tissue residence and survival^74^. They also expressed higher levels of *KIRs*, which function as inhibitory receptors^75^. In particular, the TRDV3 clusters in this atlas highly expressed *FCGR3A* (CD16), which could reflect their capacity for antibody-dependent cellular cytotoxicity (ADCC)^73^ [Figure S9G].

The expression and function of the CD4 and CD8 co-receptors have been well-studied in MAIT cells, where they can enhance the activation of the MAIT TCR^76^. However, for *γ*δ T cells, co-receptor expression has been described^77^ and identified in specific disease conditions^78^, but their role in *γ*δ T cell activation has not been well-studied nor associated with the TRDV classification. To characterise co-receptor expression on *γ*δT cell subtypes, we added another layer of co-receptor annotation on top of the *TRDV* gene usage [Figure 4A, C, S9G]. We observed that more than half of the *γ*δ T cell clusters (10 out of 18) were DN for both *CD4* and *CD8A/B*. The remainder expressed either *CD8A* only or both *CD8A* and *CD8B*, and no *CD4*^+^ *γ*δ T cell clusters were identified. Interestingly, gene expression of the CD8 heterodimer was mostly found on the *TRDV1* and *TRDV3* subtypes, while the *TRDV2/GV9* clusters were mostly DN (eight out of ten) with only 2 clusters weakly expressing *CD8A* [Figure S9G].

Since the TRDV classification was done based on gene expression, we also examined the accompanying TCR sequencing of TRDV and TRGV chains to validate the annotations. We extracted all annotated *γ*δ T cells with paired *γ*δ TCR sequencing, obtaining 4783 cells [Figure 4G], and then split them by *CD8A/B* gene expression to examine their TCR gene usage. In congruence with the gene expression-based annotation, the DN (3541/4783) and *CD8A* (116/4783) cells were mostly *TRDV2/GV9*, while the *CD8AB* (1126/4783) cells were mostly *TRDV1* (979/11126) with a small proportion of *TRDV2* (69/1126)*, TRDV3* (52/1126) and other *TRDV* (26/1126) genes [Fig 4G]. These other *TRDV* gene-expressing cells consisted of TCR-sequenced, non-*TRDV1* cells which clustered together with *TRDV1*-expressing cell clusters.

This relationship between co-receptor expression and TCR gene usage could potentially be explained by how the different *γ*δ TCRs recognise antigens. The *TRDV1* subtype has been shown to recognise MHC-like proteins from the CD1 family, namely CD1c, CD1d, Annexin A2, and MHC class I chain-related protein A and B (MICA/B)^73^, and the *TRDV3* subtype also recognises antigens in a similar way^69,73^. Conversely, the *TRDV2/GV9* subtype does not recognise antigens bound to MHC or MHC-like molecules. Instead, it detects phosphoantigens bound to the butyrophilin family protein, BTN3A1^73^. Given that co-receptor engagement can enhance antigen-dependent TCR activation with non-classical MHC-like molecules for MAIT cells^76^, NKT cells^79^, and CD1b-restricted T cells^80^, we speculate that it is possible for co-receptor engagement to play a functional role in enhancing antigen-dependent activation in *TRDV1* and *TRDV3 γ*δ T cells too. Meanwhile, since *TRDV2/GV9 γ*δ T cells do not require MHC-like presentation of antigens, co-receptor expression would confer no advantage for activation and hence this subset would be DN for co-receptor molecules. Thus, using scTCRseq alongside our transcriptomic atlas to enhance interpretation, we propose a mechanistic link between CD8 co-receptor expression, *γ*δ *TRDV* gene usage and *γ*δ TCR antigen recognition and activation.

### Combined gene expression and TCR analysis reveal functional diversity in the iNKT TCR

The final category of unconventional T cells was the DN T cells, so named for their lack of both CD4 and CD8 co-receptors^81^. They have been characterised in various conditions such as inflammation, infection, autoimmune diseases and cancer, and their function and phenotype are very much context dependent^81^. In this atlas, DN T cells were defined as cells which expressed CD3 genes but did not express the CD4 or CD8 co-receptors. They also did not strongly express genes in the MAIT or *γ*δ T cell modules [Figure S3B, S3D], but otherwise displayed characteristics of mature T cells (i.e. not thymocytes). We employed this definition to distinguish them from the MAIT and *γ*δ T cell subtypes that also did not express CD4 and CD8. In total, we identified 11 clusters of DN T cells, of which four expressed some CD4 T helper subtype-like transcription factor genes and were annotated accordingly [Figure 4A, B]. One particular cluster, DN_3, stood out due to its high expression of the TCR genes *TRAV10* and *TRBV25-1*, which identified it as a potential type I NKT or invariant NKT (iNKT) cluster^82^ [Figure 4H]. These cells have been shown to recognise lipid antigens presented by the MHC-like molecule CD1d^69^. Examination of this cluster’s DEGs showed higher expression of *ZBTB16*, *RORC*, and *KLRB1* [Figure 4H], which are some of the characteristic markers of iNKTs^83^. Functional pathway enrichment analysis using Metascape also showed an activated phenotype of cytokine and inflammatory signalling [Figure S9H].

To confirm this cluster’s TCR gene usage, we next extracted the cells in DN_3 with accompanying scTCRseq (71 out of 305 cells) and analysed them. The paired variable gene usage within this cluster was indeed biased towards *TRAV10* – *TRBV25-1* (57 out of 71 cells) [Figure 4I], matching the gene expression data. Moreover, we observed that while the alpha chain was restricted to the known invariant *TRAV10*-*TRAJ18* rearrangement, the beta chain consisted of *TRBV25-1* rearranged with various *TRBJ* genes [Figure S9I]. This resulted in a CDR3⍺ sequence that was mostly identical, but a CDR3β sequence that was highly variable across cells in the cluster [Figure 4J]. The functional significance of this phenomenon has been described by Matulis and colleagues, where they used tetramer staining to show that the hypervariable CDR3β loop dictated the affinity of the iNKT TCR to CD1d regardless of the bound ligand, and therefore determined the capacity of the TCR for autoreactive activation, even with an invariant alpha chain^84^. Consequently, iNKT clones with higher overall iNKT TCR:CD1d affinity would have an intrinsically greater autoreactive potential than low affinity clones^84^. Thus, our fine clustering and annotation has allowed us to identify and characterise a small population of iNKT cells, which when analysed together with scTCRseq data, allowed us to recapitulate a phenomenon that has been described with other techniques. Although the binding affinities of our iNKT TCRs to CD1d are not available, they add to the list of known iNKT TCRs. Further validation work could make it possible to infer the affinity of each CDR3β to CD1d and thus contribute towards iNKT TCR engineering for therapy.

### Detailed annotation and characterisation reveal experimental NK phenotypes in physiological settings

Next, we focused on the ILCs for in-depth annotation and phenotyping. ILCs differ from conventional T cells in that they do not rely on antigen-specific receptors to mount an immune response. They encompass both cytotoxic ILCs or NK cells and helper ILCs^3^, and are generally categorised into Group 1 ILCs comprising NK cells and ILC1s, Group 2 ILCs consisting of ILC2s, and Group 3 ILCs with ILC3s and lymphoid tissue-inducer cells (Lti)^85^. In this atlas, we characterised NK cells and their subsets separately from helper ILCs1/2/3, referring to them as NK cells and helper ILCs respectively. Clusters were annotated with differentiation and functional phenotypes [Figure 5A, C, S10A], as indicated by the expression of defined marker genes [Table S4, Methods]. We also visualised cluster-specific disease and tissue group proportions [Figure 5A, S10B-E] and cluster enrichment in each group was computed using Fisher’s exact test [Figure 5D, E].

**Figure 5.**
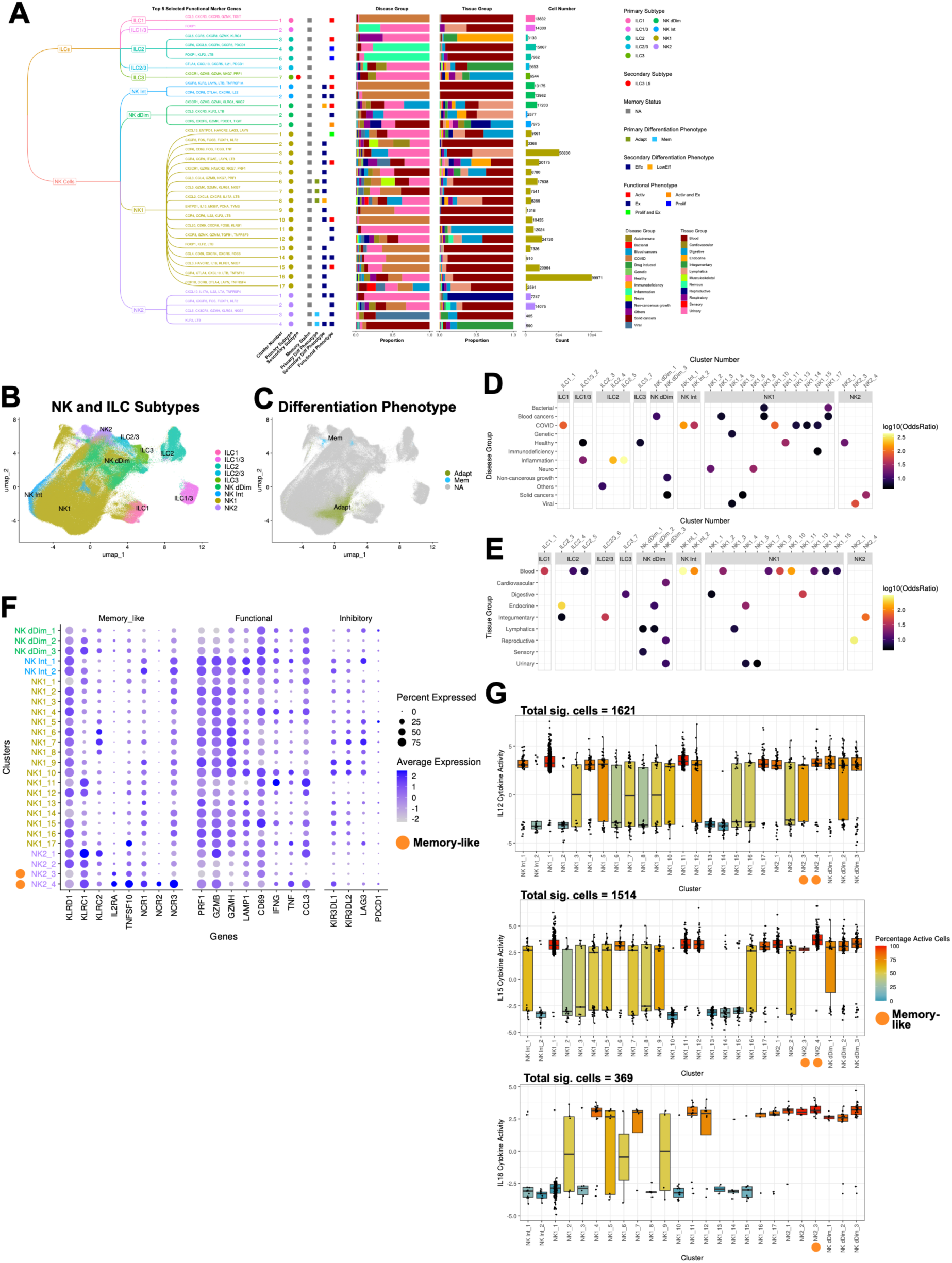
Innate cell compartment. (**A**) Tree diagram showing Natural Killer (NK) cells and helper Innate Lymphoid Cells (ILCs) and their subtypes, the clusters belonging to each subtype (named by number) and the top five selected functional marker genes of each cluster. The primary and/or secondary subtype annotations of each cluster are indicated by the coloured circles, while the primary and/or secondary differentiation phenotype and functional phenotypes are indicated by coloured squares. Stacked bar plots show the proportion of cells of each cluster coming from the various disease and tissue groups, with the cell numbers plotted in bar plots beside, coloured by subtype. (**B**) UMAP of NK and ILC subtypes, coloured by subtype. (**C**) UMAP of NK and ILC subtypes, coloured by differentiation phenotype. (**D**) and (**E**) Dot plot heatmaps showing NK and ILC clusters significantly enriched in the various disease and tissue groups respectively [P < 0.05 after Bonferroni correction and Odds Ratio (OR) > 4]. (**F**) Dot plot showing expression levels of selected genes for only the NK cell clusters, grouped and coloured by subtype. (**G**) Box plots showing the responsiveness of NK cell clusters to IL12, IL15 and IL18 as predicted by CytoSig. The gradient colour of the box depicts the percentage of cytokine-active cells in the cluster.

Helper ILCs have diverse roles in infection, inflammation, autoimmune disease, cancer, and even pregnancy and tissue repair^3,85^. They are generally classified into three subtypes which mirror the Th1, Th2 and Th17 CD4^+^ subtypes, based on similarities in transcription factor and cytokine expression profiles^3^. In our atlas, we identified seven clusters of helper ILCs according to the main transcription factors they express [Figure 5A, B]. Two of these clusters displayed an intermediate phenotype and were labelled as ILC1/3 and ILC2/3. This could reflect their functional flexibility where they can convert between subsets according to their environment and the cytokine stimuli to which they have been exposed^85^.

NK cells have been more extensively studied and are generally classified into two main groups based on their expression of CD56 (*NCAM1*) and CD16 (*FCGR3A*). The more cytotoxic NK1 is defined as CD56^dim^CD16^+^ and forms the majority of NK cells (∼90%), while the smaller population of NK2 (∼10%) is CD56^bright^CD16^-^ and is less cytotoxic but more capable of cytokine release^86,87^. In this atlas, we identified 17 clusters of NK1, four clusters of NK2, and five clusters with two intermediate phenotypes [Figure 5A, B]. The first intermediate phenotype consisting of two clusters expressed both *NCAM1* and *FCGR3A* (CD56^+^CD16^+^), which we labelled as NK Intermediate (Int) [Figure S6A]. It has been suggested that these CD56^+^CD16^+^ NK cells might represent a functional intermediate in the transition from NK2 (CD56^bright^CD16^-^) to NK1 (CD56^dim^CD16^+^), with the acquisition of CD16 as a marker of NK cell maturity^88^. The other intermediate phenotype with three clusters expressed low amounts of both NCAM1 and FCGR3A, and were labelled as NK double-dim (dDim) clusters. They are thought to differentiate from NK1 (CD56^dim^CD16^+^) cells after activation, upon which CD16 is shed from the NK cell surface by metalloprotease action^89^. Hence, these CD56^dim^CD16^-^ NK cells are probably the most mature subset of NK cells. Their numbers have been observed to increase in melanoma tumours and are correlated with improved outcome in patients receiving a cancer vaccine^90^.

While NK cells are supposed to be innate responders in the immune system, certain NK cell subtypes show memory-like characteristics where they display enhanced response upon rechallenge^91^. These are the cytokine-induced memory-like NK cells and the human cytomegalovirus (HCMV) responsive adaptive-like NK cells^91^. Cytokine-induced memory-like NK cells differentiate in response to IL12, IL15, and IL18 activation *in vitro*, and show enhanced IFNγ production after restimulation with the same cytokines^91^. They are described to express CD94 (*KLRD1*), NKG2A (*KLRC1*), NKG2C (*KLRC2*), TRAIL (*TNFSF10*), and CD25 (*IL2RA*)^91,92^. In this atlas, we identified two clusters of cytokine-induced memory-like NK cells with the gene signature above [Figure 5A, F]. The first, NK2_3, originated mainly from blood samples of viral diseases such as dengue, while the other, NK2_4, were primarily from tissue samples of uveal melanoma [Figure 5D, E, S10D, E]. Of these two clusters, NK2_4 potentially represented a cluster that had undergone activation, due to its higher expression of *NCR1* (NKp46), *NCR2* (NKp44), and *NCR3* (NKp30), especially *NCR2* which is upregulated after cytokine activation^93,94^ [Figure 5F]. Furthermore, it had higher expression of the cytotoxicity-related genes, *PRF1*, *GZMB*, *IFNG*, and *TNF* [Figure 5F].

We followed up with an *in silico* assessment of response to cytokine stimulation with the cytokine signalling analyser (CytoSig) tool, which uses transcriptomic profiles to predict the responsiveness of cells to cytokine signals^95^. We obtained predictions for IL12, IL15, and IL18 responsiveness for both clusters, except for NK2_4 with IL18 due to insufficient cell numbers. There were significant predictions for 1621 cells for IL12, 1514 cells for IL15, and 369 cells for IL18. Consistent with their annotation as cytokine-induced memory-like NK cells, both clusters displayed higher responses to IL12, IL15, and IL18 relative to most of the other NK clusters, as seen from the percentage of active cells [Figure 5G]. In particular, NK2_3 showed a high response to IL18, while NK2_4 showed a high response to IL15 [Figure 5G]. Thus, using CytoSig, we have conducted *in silico* validation of the cytokine-induced memory-like phenotype of two clusters of NK cells. This phenotype is generally induced in NK cells *ex vivo* as part of a therapeutic procedure or clinical trial^96,97^. Here, we have identified them in patients who have not undergone such NK cell therapy, which suggests that this phenotype could arise as part of normal human physiology under disease conditions.

### Cross-study comparisons and analyses of immature cells enable identification and characterisation of diseased cell types and states

The final group of cells we characterised in this atlas were the immature cells. Here, we annotated three main lineages of immature cells based on their marker genes and tissue of origin, namely the thymocytes, HSCs, and stem-like cells [Figure 6A, B, Table S4, Methods]. As with the other cell types, the disease and tissue group proportions were visualised by cluster [Figure 6A, S11B-E] and cluster enrichment in each group was computed using Fisher’s exact test [Figure 6C, D].

**Figure 6.**
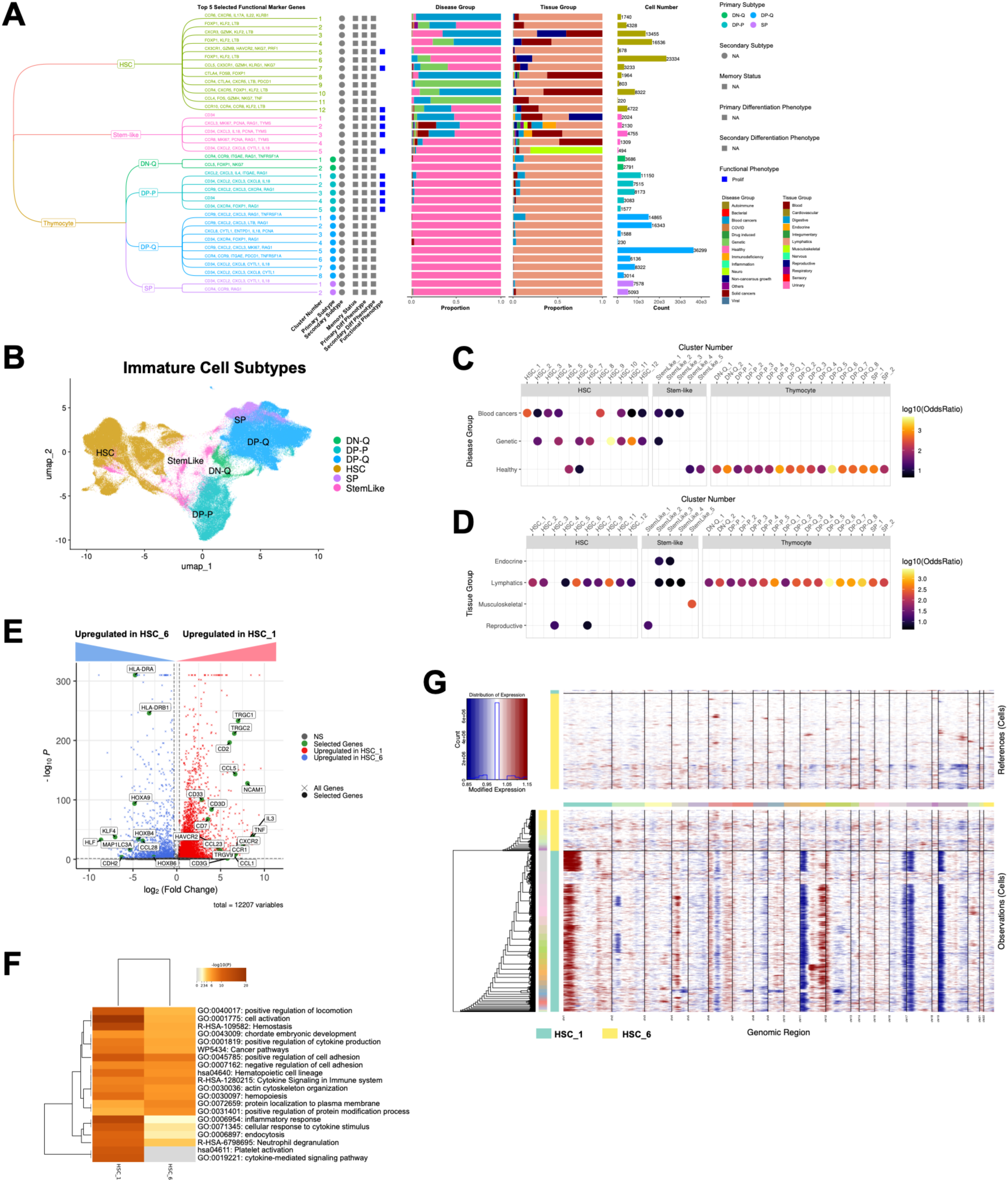
Immature cell compartment. (**A**) Tree diagram showing Hematopoietic Stem Cells (HSCs), Thymocytes and Stem-like cells and their subtypes, the clusters belonging to each subtype (named by number) and the top five selected functional marker genes of each cluster. The primary and/or secondary subtype annotations of each cluster are indicated by the coloured circles, while the functional phenotypes are indicated by coloured squares. Stacked bar plots show the proportion of cells of each cluster coming from the various disease and tissue groups, with the cell numbers plotted in bar plots beside, coloured by subtype. (**B**) UMAP of Immature cell subtypes, coloured by subtype. (**C**) and (**D**) Dot plot heatmaps showing Immature cell clusters significantly enriched in the various disease and tissue groups respectively [P < 0.05 after Bonferroni correction and Odds Ratio (OR) > 4]. (**E**) Volcano plot showing the DEGs of HSC_1 vs HSC_6. (**F**) Top 20 enriched functional pathways in HSC_1 and HSC_6 generated by Metascape, based on differentially-expressed genes (DEGs) calculated between HSC_1 vs HSC_6. (**G**) Heatmap generated by inferCNV showing chromosomal abberations in HSC_1 and HSC_6. The top panel depicts healthy cells from both clusters while the bottom panel depicts diseased cells.

The thymocyte clusters generally expressed the *RAG1* and *RAG2* genes at a higher level than other clusters [Figure S11F] and were also identifiable based on their tissue of origin, the thymus [Figure 6A, S11E]. We further split them into subsets based on their developmental stages, namely Double Negative Early (DN-E), DN Proliferative (DN-P), DN Quiescent (DN-Q), Double Positive Proliferative (DP-P), DP Quiescent (DP-Q) and Single Positive (SP) [Figure 6A, S11F], as described previously by others^65^. Of note, the DN-E and DN-P stages, representing the earliest stages of T cell development within the thymus, were not observed in this atlas in thymic tissue derived samples. Instead, we found DN-E and DN-P gene expression patterns in cells from samples of bone marrow, from which these precursors originate [Figure S11E, F]. Since the tissue of origin did not warrant a thymocyte annotation, these clusters were annotated as HSCs instead of thymocytes. In most cases, we annotated HSC clusters based on their expression of *CD34* [Figure S11F] and bone marrow tissue of origin [Figure S11E], with a few HSC clusters that came from blood samples [Figure 6A, S11E]. The final category of immature cells clustered in between the HSCs and thymocytes, but did not display any of the defining marker genes of either, and hence, were placed in a separate category on their own called Stem-like cells.

Among the HSC clusters, we observed that some displayed almost exclusive membership to a particular disease group. Here we selected HSC_1, significantly enriched in blood cancers and primarily derived from acute leukaemia of mixed phenotype (MPAL) samples, for deeper characterisation by comparing against healthy cells [Figure S11D]. For the healthy comparison, we employed HSC_6 whose cells came mostly from healthy samples [Figure S11D]. While HSC_5 also mostly came from healthy samples, it was not chosen because it had a proliferative phenotype that might have added a confounding factor to the comparison. DEG analysis between HSC_1 and HSC 6 showed that the healthy HSC_6 cluster had higher expression of genes related to stem cell self-renewal and maintenance (*CDH2*, *HLF*, *HOXA9*, *HOXB4* and *HOXB6*)^98^, and quiescence (*KLF4* and *MAP1LC3A*)^98,99^ [Figure 6E, Table S7]. In contrast, the MPAL derived HSC_1 expressed many more cytokines and receptors [Figure 6E]. Functional pathway enrichment analysis using Metascape corroborated this, showing a more inflammatory and secretory phenotype for HSC_1 [Figure 6F]. Further examination of the DEGs between the two clusters revealed that HSC_1 had a higher expression of the lineage genes *CD2*, *CD7* and *NCAM1* (CD56), while the *HLA-DR* genes were downregulated [Figure 6E]. This suggested that HSC_1 might be a cluster of HSCs that had undergone malignant transformation^100^. This was further supported by the expression of *HAVCR2* (TIM3) and *TNF* [Figure 6E], both of which are upregulated in leukemic stem cells^101,102^.

To see if HSC_1 was indeed a malignant cluster, we used inferCNV^103^ to estimate the copy number variation (CNV) of cells in both HSC_1 and HSC_6, using the cells from healthy samples from both clusters as the healthy reference for computation. Indeed, the diseased cells from HSC_1 harboured many more chromosomal aberrations than those from HSC_6, especially in chromosomes 1, 11, 12, 16, 18, and 19 [Figure 6G]. Finally, given that HSC_1 originated largely from MPAL [Figure S11D], we examined the lineage genes to see if the cells originated from bilineal or biphenotypic MPAL^104^. We found HSC_1 to highly express *CD3* and *TRGC/TRGV* compared to HSC_6, indicating a T-lineage leukaemia [Figure 6E]^104^. This implied a bilineal MPAL origin where malignant cells heterogeneously expressed markers of two different lineages as opposed to a biphenotypic MPAL where different lineage markers are expressed on the same cells^104^. However, there is no matching cluster from the myeloid lineage available, likely due to its removal in the filtering step of myeloid cell removal during atlas construction. In addition, we noted that HSC_1 was a relatively small cluster of cells (1740 cells), which shows that our clustering and annotation is sensitive enough to detect and capture small but distinct cell types. Finally, tracing HSC_1 back to the main project of origin (GSE139369), we found that in contrast to the original study, which had difficulty distinguishing the various mixed-phenotype leukaemia cells from each other, our comparison with a separate healthy HSC cluster enabled us to identify the T-lineage phenotype clearly. Thus, this atlas provided healthy and diseased cells from different studies for comparison, which enabled us to identify, analyse and validate with *in silico* tools, a small malignant leukemic stem cell cluster originating from MPAL.

### Validation of a tumour tissue-infiltrating and immunosuppressive CD4^+^ Treg cluster with spatial transcriptomics

Finally, to show that our fine grained clustering and annotations are useful to direct further validation and discovery, we selected CD4_Treg_8 for validation, identified earlier as an immunosuppressive Treg cluster which infiltrates solid tumours of epithelial or epithelial-like phenotype. We collected publicly available spatial transcriptomic samples from each of the solid cancers which made up this cluster, BRCA^105^, BCC^106^, HCC^107^, NSCLC^108^, NPC^109^ and oral SCC (OSCC)^110^, focusing on 10X Visium samples as the most readily-available platform across all the cancer types, and did data integration across the samples [Methods]. This gave us an integrated dataset of 122 samples from 69 individuals, including some healthy lung tissue (n=4) and normal adjacent lung tissue (n =16) [Figure 7A], with a total of 221,361 spots for analysis [Figure S12A]. To identify which spots might contain CD4_Treg_8, we created a CD4_Treg_8 gene module comprising the tumour-specific signature identified earlier (*CXCL13, KLRB1, ADGRG1, BHLHE40, MYO7A, PTPN13, CSF1*) [Figure 2F] and some CD4^+^ Treg-related genes (*CD4, IKZF4, FOXP3, IL2RA, CTLA4*), and applied Seurat’s AddModuleScore function to score the expression of the CD4_Treg_8 module in each spot. We selected the top 1% of spots by score, and the top 5% of spots with the most even distribution of the module genes to ensure that the module score was not skewed by any single gene, obtaining 1741 spots with high and even expression of the CD4_Treg_8 gene module (CD4_Treg_8-high) [Figure S12B]. Analysis of the CD4_Treg_8-high spots by tissue type revealed that they were enriched in tumour tissue across all the cancers except for HCC [Figure 7B]. In contrast, no spots with CD4_Treg_8 signature were detected in healthy tissue, and only one spot was found in one normal adjacent tissue sample of NSCLC [Figure 7B]. The almost exclusive enrichment of the CD4_Treg_8 signature in tumour tissue was statistically significant (Wilcoxon rank-sum test, p=0.00015) [Figure 7C]. This validated our previous finding from the atlas that CD4_Treg_8 preferentially infiltrated tumour tissue.

**Figure 7.**
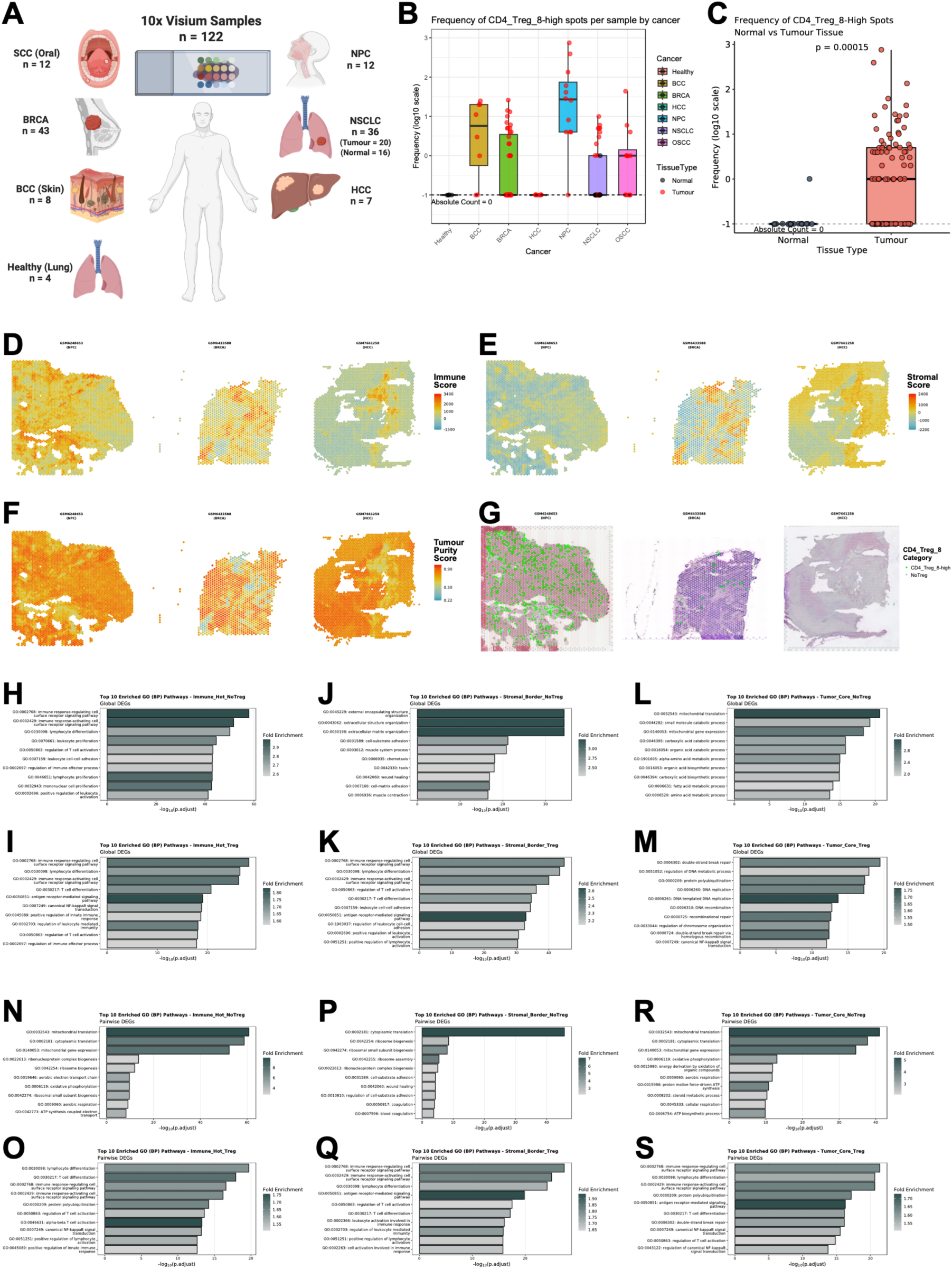
Validation of CD4_Treg_8 with spatial transcriptomics. (**A**) Schematic showing the breakdown of 10x Visium tissue samples for each of the epithelial malignancies. (**B**) Boxplots showing the frequency of CD4_Treg_8-high spots per sample, split by cancer type. The colour of the dots show the tissue type (normal vs tumour tissue) (**C**) Boxplots showing the enrichment of CD4_Treg_8-high spots in Tumour vs Normal tissue (Wilcoxon rank-sum test, p=0.00015). (**D**) Representative spatial dimension plots showing spot-level Immune Score as calculated by ESTIMATE. (**E**) Representative spatial dimension plots showing spot-level Stromal Score as calculated by ESTIMATE. (**F**) Representative spatial dimension plots showing spot-level Tumour Purity Score as calculated by ESTIMATE. (**G**) Representative spatial dimension plots showing infiltration of CD4_Treg_8. (**H-M**) Top 10 Gene Ontology (GO) biological pathways (BP) for CD4_Treg_8-high (Treg) vs No CD4_Treg_8 (NoTreg), for globally calculated DEGs (vs all other regions). (**N-S**) Top 10 GO-BP for CD4_Treg_8-high (Treg) vs No CD4_Treg_8 (NoTreg), for pairwise calculated DEGs (within each local environment: Immune_Hot, Stromal_Border and Tumour_Core). Schematic for (**A**) created with BioRender.com.

Next, we investigated the effect that CD4_Treg_8 might have on its immediate local microenvironment in tumour tissue. Focusing only on tumour tissue, we employed the Estimation of STromal and Immune cells in MAlignant Tumours using Expression data (ESTIMATE) tool^111^ to infer the fraction of stromal and immune spots in our tumour samples. This analysis returned an Immune Score, a Stromal Score, and a Tumour Purity Score derived from the former two scores for each spot, which could then be visualised on feature plots [Figures 7D-E]. Spots identified as CD4_Treg_8-high were also visualised alongside the ESTIMATE scores [Figure 7G]. We then stratified all the spots into discrete microenvironment categories based on the ESTIMATE scores and CD4_Treg_8 infiltration, generating six mutually exclusive compartments: Immune_Hot (highest relative immune cell infiltration), Stromal_Border (highest relative extracellular matrix or fibroblast presence), and Tumor_Core (highest relative malignant cell purity) for both CD4_Treg_8-infiltrated (Treg) and CD4_Treg_8-absent (NoTreg) niches.

To characterise each compartment individually, we conducted a global DEG analysis, comparing the gene expression profile of each compartment to the entire dataset [Table S8], followed by functional pathway enrichment analysis, focusing on Gene Ontology (GO) biological pathways (BP), to uncover the key functional pathways and biological mechanisms affected by the presence of CD4_Treg_8. In Immune_Hot compartments with highest immune cell infiltration, an immune cell proliferative phenotype (leukocyte, lymphocyte and mononuclear cell proliferation) was observed in the absence of CD4_Treg_8 (NoTreg) [Figure 7H], likely indicating an ongoing and active immune response. In contrast, Immune_Hot compartments with CD4_Treg_8 (Treg) did not have the same proliferative phenotype. Instead, more immune regulatory or dampening pathways were observed such as the regulation of leukocyte mediated immunity, T cell activation, and immune effector processes [Figure 7I]. Similarly, for the Stromal_Border compartment, the absence of CD4_Treg_8 showed an enrichment of processes involved in extracellular matrix organisation [Figure 7J], expected in a fibroblast-enriched niche, while the presence of CD4_Treg_8 displayed greater immune activity, including regulatory pathways and immune dampening [Figure 7K]. Finally, Tumour_Core compartments without CD4_Treg_8 were enriched in metabolic processes [Figure 7L], while the presence of CD4_Treg_8 was associated with DNA replication, repair and organisation [Figure 7M], indicating a more proliferative tumour cell phenotype and possibly a more aggressive tumour environment. We also conducted pairwise DEG comparisons within the Immune_Hot, Stromal_Border and Tumour_Core compartments, with versus without the presence of CD4_Treg_8 [Table S9]. In congruence to the global DEG analysis, in the absence of CD4_Treg_8, each compartment showed an enrichment in phenotypes related to its corresponding cell type functions, such as protein translation and respiration for immune cell activity [Figure 7N], protein translation and cell adhesion for stromal cell activity [Figure 7P], and respiratory processes for tumour cell activity [Figure 7R]. In the presence of CD4_Treg_8, all compartments uniformly demonstrated an enrichment in pathways involving immune activity, including immunosuppressive functions such as regulation of T cell activation [Figures 7O, Q, S]. We conclude therefore, that CD4_Treg_8 has an immunosuppressive effect on its immediate microenvironment. Thus, we have validated the presence of an immunosuppressive Treg cluster which selectively infiltrates tumour tissue in epithelial malignancies.

## Discussion

Here, we present Uni-TINT, a pan-disease, pan-tissue atlas of T and ILCs aimed at unifying T and ILC taxonomy. Through careful compilation of marker genes, a systematic and rigorous method of hierarchical manual annotation, and detailed metadata curation at the sample level, we provide highly granular clusters of T/ILCs, achieving the depth of annotation that individual studies can provide, and interpreting their function and phenotype in the context of disease and tissue of origin.

Within the CD4^+^ T cell compartment, we characterised nine clusters of CD4^+^ Tregs with a variety of phenotypes, including Tr1, Tfr, cytotoxic, inflammatory, autoimmune-associated, virus-infected, and solid cancer-associated Tregs. Some of these have been identified using other technologies such as flow cytometry but have not been characterised in scRNAseq-derived data. Others are novel either in terms of their phenotype or the disease context where they are found. Access to data from various studies enabled us to identify Treg clusters enriched within a disease group and yet distributed across diseases within that group, which gives rise to the exciting possibility of targeting these cell types for therapy. In particular, we characterised a solid cancer-associated Treg cluster, CD4_Treg_8, from a variety of epithelial malignancies with a highly immunosuppressive phenotype. Using the gene signature expressed by this cluster, we validated its tumour tissue-infiltrating and immunosuppressive properties in spatial transcriptomic tissue samples of the same epithelial malignancies. It is possible that in tumorigenesis or cancer progression, pathological CD4^+^ Treg phenotypes develop which are common across solid cancers. The solid cancer-associated CD4_Treg_8 represents a potential target for immunotherapy across cancers.

In a similar way, within the immature cell compartment, cross-study analysis enabled us to identify and characterise a small malignant HSC cluster by comparison to healthy HSCs. This was achieved even without cytomorphologic evaluation or flow cytometry, which are the primary means of diagnosing acute leukaemias^112^. Bilineal MPAL presents with at least two different populations of leukemic cells from the B cell, T cell or myeloid lineages in the same patient^104^. Using transcriptomic analyses and *in silico* tools, we were able to single out and phenotype malignant cells from the T lineage, but not from the other lineages because they would have been filtered away as non-T cells at the quality control step of building this atlas. Our fine-grained clustering thus facilitated the detection and characterisation of a small diseased cell type. The ability to conduct cross-study analyses thus expands both the breadth and depth of immune profiling possible, with comprehensive coverage across diseases, and sensitive detection of small discrete phenotypes.

Within the CD8^+^ and Unconventional T cell compartments, we identified subsets of cells which would normally be difficult to profile in an individual study due to their rarity. These are the *FOXP3*^+^ CD8^+^ Tregs, the *KIR*^+^ CD8^+^ Tregs and the iNKTs. Notably, we characterised *FOXP3*^+^ CD8^+^ Tregs from the thymus and iNKTs using scRNAseq for the first time. The origin of *FOXP3*^+^ CD8^+^ Tregs is disputed, with various studies arguing for or against thymic or peripheral origins^62,113^. Here, we detect them in the thymus with a transitioning phenotype, which suggests that they might indeed originate in the thymus. However, we did not detect their counterpart in the periphery and therefore were unable to compare or track their development. The iNKT subset, on the other hand, has been of particular interest due to its potential autoreactive capabilities^114^. While certain autoreactive iNKT clones have been identified, the identities of their self-antigens remain elusive^114^. Here, we demonstrate that the expression of the iNKT TCR also sets this subset apart in terms of gene expression. In combining GEx and TCR analyses, we recapitulate the phenomenon where the CDR3β loop determines the diversity of the iNKT TCR repertoire^84^, and add to the pool of known iNKT TCR sequences, which could potentially be used to screen for autoreactive function and self-antigens.

Combining scRNAseq with scTCRseq to enhance functional characterisation for *γ*δT cells, we found that co-receptor expression appeared to correlate with TRDV chain usage and classification, and that this phenomenon was potentially explainable by how different *γ*δ TCRs recognise their antigens. Various immunotherapeutic strategies are being explored which involve *γ*δT cell engineering, including Chimeric Antigen Receptor (CAR) *γ*δT cells and engineering ⍺β T cells to express a *γ*δ TCR^115^. A fuller understanding of the mechanics of co-receptor engagement and antigen recognition by the different *γ*δ TCRs might make it possible to tailor these immunotherapies to enhance or modulate the desired therapeutic effect, depending on the target antigen and *γ*δ TCR being used.

Finally, within the innate cell compartment, we detected an unconventional phenotype of NK cells, namely the cytokine-induced memory-like NK cells and validated it using *in silico* methods. Cytokine-induced memory-like NK cells have shown promise in cancer immunotherapy. The ability to generate a memory phenotype and boost IFNγ production upon restimulation raised the exciting possibility of administering such cells as treatment. Indeed, they have shown great promise in *in vitro* experiments, *in vivo* models and in clinical trials^96^, and even as the effector cell in CAR therapy^97^. Common across all these trial therapies is that these NK cells are preactivated *ex vivo* before administration into the subject. The identification of the cytokine-induced memory-like phenotype in individuals who have not undergone such therapy suggests that these cells could develop as part of normal human physiology in some disease contexts, and raises a new and exciting avenue of immunotherapy, that of targeting and reactivating these cells in patients without the need for *ex vivo* stimulation.

In summary, Uni-TINT addresses several needs faced in in-depth immune profiling, such as the lack of consensus definitions of cell type and state, the absence of a catalogue of finely annotated cell clusters to compare against, and the need for a database which comprehensively covers both disease and tissue types. We compile and refine T and ILC taxonomy to provide a detailed and consistent reference for researchers to use. Using this atlas of 3.47 million transcriptomes across various disease and tissue types, we have shown how fine-grained annotation alongside disease and tissue information greatly enriches our understanding of the immune contexture and interpretation of immune function. We also demonstrate how Uni-TINT, when supplemented by antigen receptor sequencing and various *in silico* tools, can provide a primary layer of validation when screening for cell types or targets of interest. Finally, we validated a tumour tissue-infiltrating CD4 Treg cluster, providing confidence for further validation of other clusters of interest. Thus, Uni-TINT is not just a reference map, but also serves as a platform for discovery and downstream validation, and for application to more advanced computational approaches. Uni-TINT will be available on DISCO, and the annotations will be incorporated into our cell type annotation tool CELLiD. This study has focused mainly on disease and tissue effects on immune cell function. Further curation and investigation are needed for other important sample-level variables such as age, gender and HLA type for an even more comprehensive understanding of the immune contexture.

## Supporting information

List of Abbreviations

## Data and code availability

This study analysed publicly available datasets obtained from the Gene Expression Omnibus (GEO), Sequence Read Archive (SRA), and other public repositories. Accession numbers for all datasets analysed are provided in the Supplementary Information. No new sequencing or other standardized datasets were generated in this study. The datasets supporting this study are available on Zenodo at DOI: [10.5281/zenodo.21667682 - PART ONE], DOI: [10.5281/zenodo.21711927 - PART TWO], and DOI: [10.5281/zenodo.21716235 – PART THREE]. The source code, custom analysis scripts, and pipeline instructions used to generate the Uni-TINT atlas are available at https://github.com/JinmiaoChenLab/Uni-TINT. These repositories will become publicly accessible upon peer-reviewed publication. An interactive version of Uni-TINT is available as a ShinyCell2^116^ app at https://disco.bii.a-star.edu.sg/Uni-TINT.

## Acknowledgements

We would like to thank Professor Nicholas Gascoigne for his valuable insights, and members of both CJM and NRG groups for their useful discussions. We thank Dr Mengwei Li for his work in building the DISCO database, which is the backbone of this atlas.

Funding was provided by the National Research Foundation Investigatorship (NRFI), Singapore (NRF-NRFI11-2026-0014), the Ministry of Education Academic Research Fund Tier 2, Singapore (MOE-T2EP30125-0017) and the NMRC Open Fund Large Collaborative Grant, Singapore (MOH-001575-00) grants.

## Contributions

J.C. conceptualised and supervised the project. N.R.J.G. supervised the project. S.W.J.C. built the atlas and did the analyses. M.L. provided support via DISCO. S.W.J.C., J.C. and K.S.A. wrote the manuscript.

## Methods

### Data collection and computational extraction of T cells and Innate Lymphoid Cells (ILCs)

The atlas was constructed using publicly available scRNAseq data which were previously processed with a standardised pipeline and made available on DISCO^30^. In the processing pipeline, QC was performed and the raw reads were mapped onto the human reference genome assembly hg38. For sample selection, the following criteria was employed. First, only samples sequenced with the 10x Genomics platform were retained to reduce technical variation in the data. Second, the following types of samples were excluded, samples with genetic modifications, without proper metadata, and sorted populations of non-T and ILCs. The resulting list of 1869 samples from 194 studies is provided in [Table S1].

The first step of atlas construction involved T and ILC identification and extraction by coarse-grained clustering and annotation. As the assembled data was too large for efficient handling as a single dataset on the R platform, the samples divided up by pseudobulking individual samples and clustering them to obtain 23 clusters. The samples within each cluster were then processed and integrated together. The Seurat v4^117^ and scDblFinder^118^ libraries were used for quality control, doublet removal, and normalisation of individual samples. Samples of each cluster were then integrated using the FastIntegration R package^30,119^. This was followed by PCA and clustering to obtain clusters of major cell types, which were automatically annotated at the cluster level with using CELLiD^29,30^. Based on CelliD’s annotation, cell barcodes of the T cells, ILCs, and hematopoietic stem cells (HSCs) were extracted. The HSCs were included to enable analysis of immature cells.

### Integration, clustering, manual annotation and refinement of data

For the next step in constructing the atlas, the selected T/ILC cell barcodes were used to extract the original raw gene expression. The extracted data was integrated using the FastIntegration package^30,119^ using sample IDs as batch labels and 2000 highly variable genes selected via Seurat’s FindVariableFeatures function using the vst method. [Figure S1A]. To refine and annotate the atlas, two different stages of iterative clustering and annotation were performed.

In the first stage, the goal was to identify the main cell types (CD4 T, CD8 T, unconventional T, ILCs with NK, and immature) and remove any remaining non-T/ILCs present. At each iteration, a stable clustering was selected. The corresponding clusters were downsampled to 2000 cells per cluster for differentially expressed genes (DEGs) computation with Seurat’s FindMarkers function by comparing each cluster against the rest [Table S5]. Each cluster was then manually annotated by searching the DEG lists for cell type markers [Table S4]. Identified clusters of non-T/ILCs such as B cells and myeloid cells were marked and removed. This process was repeated with progressively finer clustering resolutions to achieve pure T/ILCs clusters [Figure 1C].

After removing contaminating cells, the dataset was split by the major cell types, i.e. T cells, NK cells, ILCs, and others, for the second stage of clustering and annotation. Within each main cell type, a similar process was employed [Figure 1C], namely clustering and manual annotation by computing for DEGs against the entire atlas dataset [Table S5] and cell type gene set scoring using Seurat’s AddModuleScore function [Table S4]. This process was repeated until sufficiently pure clusters with finer cell subtype annotations were achieved.

All clustering performed on the atlas employed FastPG^120^. To determine if clustering was sufficiently fine, four methods were employed concurrently. 1) FastPG^120^ clustering was conducted for different values of k, and the results were visualised using the R package Clustree^121^. Clustree provides an easy visualisation of the stability of a cluster across different clustering resolutions. 2) Clusters were visualised on UMAP to see if they could be split into subclusters. 3) The DEGs of each cluster were calculated and examined for a reasonable phenotype. 4) Cluster purity was calculated using the ROGUE score^122^ to ensure that sufficient clustering had been achieved to give homogeneity within a cluster and across the disease groups [Figure S1B].

### Detailed manual annotation procedure and rationale

The following manual annotation procedure was performed on each of the 207 clusters individually. Seurat’s AddModuleScore function was performed on uncorrected data and key marker genes were also examined from each cluster’s DEG to make the final decision for the cell type annotation. The module’s score and key marker genes were visualized on violin plots. Genes which appeared in the DEG of a cluster [Table S5] were considered as highly expressed for that cluster. A module’s score for a cluster would also be considered high if it was higher relative to the other clusters. Key marker genes used to analyse the DEG for each cluster are in Table S4. Genes used in each module are also listed in Table S4, and are stated in each section describing the annotation process below.

#### Identifying the T cells, NK cells, and ILCs

The T, NK, and ILC clusters were first broadly separated using the CD3 (*CD3E*, *CD3D*, *CD3G*) and NK (*KLRD1*, *KLRF1*, *NKG7*, *XCL1*, *XCL2*) gene modules [Figure S2A]. Clusters which highly expressed all CD3 genes and had a high CD3 but low NK signature score were classified as T cells. Conversely, clusters which highly expressed the NK genes and had a high NK but low CD3 score were classified as NK cells. Clusters were labelled as ILCs if they did not show highly expressed CD3 or NK genes and had a low score for both CD3 and NK modules. Each major cell lineage’s clusters were then analysed separately for finer subtyping, as well as differentiation and functional phenotyping.

#### T cell typing and phenotyping

The T cell clusters were divided into the following main lineages, conventional CD4^+^ and CD8^+^ T cells, thymocytes, and unconventional double negative (DN), *γ*δT, mucosal-associated invariant T (MAIT), and MAIT-like cells [Fig S3A, Table S4]. At the first pass, clusters were labelled based on their CD4 and CD8 gene expression. Clusters which highly expressed *CD4* and not *CD8A* or *CD8B* were classified as CD4 T cells. Clusters which highly expressed both *CD8A* and *CD8B* were classified as CD8⍺β T cells, while those which only expressed *CD8A* were labelled as such. Clusters lacking CD4 and CD8 expression were considered DN T cells, while those with *CD4*, *CD8A*, and *CD8B* expression were labelled as double positive (DP) T cells and with thymus as tissue of origin and RAG expression, labelled as thymocytes and considered separately. As *CD4*, *CD8A*, and *CD8B* expression is not limited to the conventional subsets, these labels were also retained for the unconventional T cell clusters.

Thereafter, cell subtypes were assigned based on markers in the cluster DEGs and gene module scores, and up to one primary and one secondary subtype could be assigned. If a cluster scored highly and highly expressed most of the key genes, it would be labelled with first-level confidence as the primary subtype. If the cluster lacked either one aspect, i.e. low score but high in DEG or vice versa (which occurred more frequently), it would be labelled with second-level confidence as the secondary subtype. However, there were also clusters that displayed a mixed phenotype in terms of DEGs and/or module scores despite attempts to refine them. Such clusters were annotated with a primary subtype with higher confidence and a secondary subtype with lower confidence. Finally, clusters without key subset marker genes were labelled as NoSubset and were often naïve cells. We note that the subtyping process of CD4^+^ T cells was separate from all other T cells as some of the subsets’ markers were common to other cell types and would thus affect the computed module score.

For the conventional CD4 subsets, the following gene modules were employed: T helper type (Th) 1 (*IFNG*, *TBX21*), Th2 (*GATA3*, *PTGDR2*), Th17 (*IL17A*, *RORC*, *RUNX1*), Th22 (*IL22*, *AHR*), T regulatory (Tregs) (*FOXP3*, *IL10*), and T follicular helper (TFH) (*BCL6*, *CXCR5*, *ICOS*). For Th9 (*SPI1, IL9*), no cluster could clearly be identified [Fig S5A, Table S4]. For the CD8 cytotoxic T (Tc) cells, similar subsets were assigned, as they have been shown to have subsets that corresponded to the CD4 subsets^27^. The gene modules employed were also similar, Tc1 (*TBX21*, *EOMES*), Tc2 (*GATA3*, *PTGDR2*), Tc9 (*IRF4*, *STAT6*, *IL9*), Tc17 (*RORC*, *IL17A*, *KLRB1*), Tc22 (*AHR*, *IL22*, *STAT1*), T follicular cytotoxic (Tfc) (*CXCR5*, *BCL6*, *CD40LG*), killer cell immunoglobulin like receptor (KIR) expressing cells (*KIR3DL1*, *KIR2DL1*, *EOMES*), and CD8 Tregs (*FOXP3*, *IL10*) [Figure S5B, Table S4].

To annotate the *γ*δT cell clusters, three gene module scores for the constant (*TRDC*, *TRGC1*, *TRGC2*), variable delta (*TRDV1*, *TRDV2*, *TRDV3*) and variable gamma (*TRGV2*, *TRGV3*, *TRGV4*, *TRGV5*, *TRGV8*, *TRGV9*) were calculated [Figure S3B, Table S4]. A cluster would only be labelled as *γ*δT if it had a high score for two or more modules and had highly expressed gamma and delta chain genes as DEGs. For subtyping, the *γ*δT clusters were classified based on their TRDV chain expression^73^; each *γ*δT cluster was labelled as TRDV1, TRDV2, or TRDV3 based on the highest expressing chain as observed in their DEG list and in the gene expression violin plots [Figure S3C, Table S4].

For MAIT cells, the MAIT gene module (*TRAV1-2*, *KLRB1*, *SLC4A10*) was used [Figure S3D, Table S4]. A cluster would be labelled as MAIT cells if it scored highly for the module and highly expressed key MAIT genes, especially *TRAV1-2*. If a cluster scored highly on the MAIT module but did not express TRAV1-2 in its DEG, it was considered MAIT-like.

Finally, thymocyte clusters were manually annotated based on *RAG1* and/or *RAG2* in the cluster DEGs [Table S5] and confirmed with the thymus being their tissue of origin, [Figure S11E].

#### T cell memory and differentiation

All T cell clusters were also split into a first-pass memory and non-memory classification using a memory differentiation gene module (*S100A4* and *GPR183*) [Figure S4A, Table S4]. Clusters with high relative expression of the module were classified as memory, while those with low expression as non-memory. This memory status was further verified when assigning the differentiation phenotype of each cluster.

After memory phenotype assignment, the T cell clusters were separated by the memory status for determining their differentiation phenotype. This division was due to certain markers (e.g. *CCR7* and *SELL*) being expressed by certain phenotypes from either group (e.g. central memory and naïve cells). To determine the primary differentiation phenotype of memory type cell clusters, central memory (CM), effector memory (EM), and tissue resident memory (TRM), the following gene modules were used. The CM gene module contained *CCR7*, *SELL*, and *CD27*, EM used *TBX21*, *CX3CR1*, *CXCR3*, *CXCR4*, *FCGR3A*, *KLRD1*, and *GZMK*, and the TRM module was defined by *ITGA1*, *ITGAE*, *PRDM1*, and *ZNF683* [Figure S4B, Table S4]. For non-memory clusters, the following gene modules were used. The naïve gene module contained *TCF7, LEF1*, and *CD27*, the secondary lymphoid module had *CCR7* and *SELL*, and the effector set consisted of *CX3CR1, IFNG, PRF1, TNF, IL4*, and *GZMB* [Figure S4C, Table S4].

To ascertain the state of terminal differentiation, all T cell clusters were also assigned a secondary differentiation phenotype based on the terminal differentiation (TEMRA) phenotype with the following gene module, *KLRG1*, *S1PR5*, and *B3GAT1* [Figure S4B, C, Table S4]. Finally, the exhaustion functional phenotype was assigned based on the genes *CTLA4*, *PDCD1*, *TIGIT*, *ENTPD1*, *LAYN*, *HAVCR2*, *CXCL13*, *TNFRSF9*, and *LAG3* [Figure S4B, C, Table S4]. These module scores, together with additional selected genes, were used to manually annotate the phenotypes of each cluster.

#### NK cell and helper ILC subtyping and phenotyping

Unlike the T cells, NK cell clusters were assigned only one (primary) subtype based on both their DEGs and their gene module scores. To differentiate between NK1 and NK2, the respective gene modules were used, NK1 (*FCGR3A*, *KIR3DL1*, *KIR3DL2*) and NK2 (*NCAM1*, *SELL*, *IL2RA*, *IL18*, *GZMK*), and visualisation of key marker expression, *NCAM1* (CD56) and *FCGR3A* (CD16) [Figure S6A, Table S4]. For clusters which showed expression of both *NCAM1* and *FCGR3A*, they were annotated as NK_Int (intermediate), and clusters with low expression of both were annotated as NK_dDim (double dim) [Figure S6A, Table S4].

Some unconventional NK cell types can display memory-like or adaptive-like phenotypes in certain contexts^91,92^. Therefore, where appropriate, NK clusters were assigned the primary differentiation phenotypes cytokine induced memory-like (*TNFSF10*, *IL2RA*) and adaptive (*KLRC2*, *B3GAT1*) [Figure S6B, Table S4]. Additionally, a secondary effector phenotype was assigned based on effector genes (*LAMP1*, *PRF1*, *GZMB*), and functional phenotype of exhaustion based on exhaustion or inhibitory markers (*PDCD1*, *KLRC1*) [Figure S6C, Table S4].

For the helper ILCs, three subtypes were annotated, ILC1 (*TBX21, IFNG, TNF, PRF1*, *IKZF3*), ILC2 (*GATA3, PTGDR2*, *IL1RL1*), and ILC3 (*RORC, AHR, IL23R*, *CCL20*) [Figure S6D, Table S4]. ILC3 were further annotated with a secondary subtype called the lymphoid tissue inducer (Lti) (*ID2*, *TNFSF11*, *LTB*). There were also clusters which showed a mix of marker genes and thus were labelled as intermediates of the subtypes with those markers [Figure S6D].

#### Immature cells subtyping and phenotyping

Based on their developmental stages, thymocytes can be categorised into DN early (E), DN proliferating (P), DN quiescent (Q), DP (P), DP (Q), and single positive (SP) [Table S4], as previously described by others^65^. In the manual annotation of thymocyte clusters, the DN (E) and DN (P) stages were not found. Instead, their phenotype was observed in some HSC clusters, which can be thymic progenitors [Figure S11F]. The HSC clusters were annotated based on their expression of *CD34* and having bone marrow as tissue of origin. Finally, some clusters displayed an immature phenotype but expressed incomplete or indeterminate combinations of key marker genes, which were labelled as stem-like cells.

#### Activation and proliferation phenotyping

The assignment of activation and proliferation functional phenotypes were based on the gene expression and scoring with their respective gene modules, activation (*CD69*, *FOS*, *FOSB*) and proliferation (*MKI67*, *TYMS*, *PCNA*) [Figure S2B, Table S4]. For T cells, *FOS* and *FOSB* are indicative of TCR signalling, hence a combination of any two of the three markers was taken as evidence of activation. For NK and ILCs, only CD69 was taken as the main activating marker, without which a cluster would not be considered as activated [Table S4].

For proliferation status, upregulation of two or more proliferation genes in the DEGs together with a high module score was required to label a cluster as proliferating. If only one proliferation gene was upregulated, but the module score was low, it was not considered as a proliferative cluster. If one gene was upregulated together with a high module score, the cluster was considered as proliferating. If two genes were upregulated while the module score was low, it was not considered as proliferating.

### Metadata curation

As part of the atlas construction, the sample metadata was harmonised across studies as much as possible. To achieve this, the source publications were manually searched for all relevant metadata. This included primary disease, disease subtype, disease stage, tissue of origin, age, treatments received, time point of sample collection, cell sorting definitions, gender and sample type. These variables were harmonised and further grouped into larger categories for ease of analysis, such as disease, age, and tissue groups [Figure 1B].

### scTCRseq data collection and processing

Available scTCRseq data of samples in the transcriptomic atlas were downloaded and processed. As the atlas was restricted to 10x Genomics Chromium sequenced samples, the scTCR data were also acquired using the same platform. Where available, the raw reads were downloaded and processed with Cellranger VDJ (version 7.1.0). Otherwise, available processed files were utilised. All processed TCR information were harmonised into a common format of clonotype, chain, and gene information. For downstream integrated analysis with the transcriptomic atlas, the TCR profiling information were matched via cell barcodes [Figure 1B].

### HLA typing with arcasHLA

The HLA type of each sample in the atlas was inferred *in silico* using arcasHLA^31^ and the IMGT/HLA version 3.34.0 database. From the sorted sample BAM files, reads that mapped to chromosome 6 as well as unmapped reads were extracted. The extracted reads were then used to predict the most likely genotype without the partial alleles option. The output contained two predicted alleles for each HLA locus, and this information was retained for homozygous predictions; the full predictions were incorporated into the atlas metadata for downstream analyses. When computing allele frequencies, all alleles per sample were counted once, even for homozygous alleles.

### Fisher’s Exact Test for enrichment

To test for enrichment of a cluster in a group, Fisher’s exact test was used. Contingency tables were first constructed for each cluster versus each group. Contingency tables with zero values in positions a or d of the table (i.e. both present and/or absent in both cluster and group) were excluded from analysis because these would give an odds ratio (OR) of 0 (fully negative association). The test statistic was calculated using the fisher.test function in R. Multiple test correction was done using the p.adjust base R function with the Bonferroni method option, and the test results were filtered by an adjusted p-value < 0.05 and OR > 4. This would mean that the statistically significant results were 4 times more likely to have cells from that cluster in that group.

### Functional pathway enrichment analysis using Metascape, scMetabolism and clusterProfiler

Functional pathway enrichment analysis was performed at the Metascape^57^ website with the Express Analysis function. For all Metascape analyses, the top 1000 DEGs for each cluster calculated against all other cells were used as input, except for the analysis of HSC_1 vs HSC_6 in Figure 6C, where the top 1000 DEGs calculated between the two clusters was used. Heatmap plots of the enriched functional pathways were automatically generated by Metascape. The clusterProfiler R package^123^ was used for Gene Ontology (GO) biological pathway (BP) specific enrichment analyses, especially for the analyses in Figures 7H-S.

For analysing cellular metabolism, the scMetabolism package^58^ in R was employed. For the CD4 Treg analysis, all nine Treg clusters were downsampled to 2000 cells per cluster and the Seurat object with their uncorrected gene expression values were used as input. The default options, VISION method, use of imputation, and KEGG metabolism type were used. The computed metabolic pathways were grouped according to broad categories and selected pathways that showed meaningful differences between the clusters were plotted using ggplot2.

### Predicting cytokine responsiveness with CytoSig

The responsiveness of NK cell clusters to cytokine signalling was calculated using the CytoSig prediction model in Python^95^. The log-normalised counts for all NK clusters (downsampled to 2000 cells per cluster) were extracted, and CytoSig was run with the default parameters except for the use of the expanded signature option (-s 2) to include a more comprehensive coverage of cytokines. Multiple test correction was done using the p.adjust base R function with the Benjamini-Hochberg (BH) method option to give the false discovery rate (FDR). The z-scores for all cells with FDR < 0.05 were extracted for visualisation. A more positive z-score indicates greater responsiveness to the cytokine. For the NK clusters, responsiveness to IL12, IL15, and IL18 were analysed.

### Chromosomal aberration analysis with inferCNV

The inferCNV R package^103^ was used to analyse the potentially malignant HSC clusters. The raw uncorrected gene expression values of HSC_1 and HSC_6 (2000 cells per cluster) were used as input, and the annotated healthy cells from both clusters were used as the healthy reference. Default parameters were used, with a cutoff of 0.1 as recommended for 10x Genomics data.

### 10x Visium spatial transcriptomics data processing and integration

The epithelial malignancy spatial transcriptomic dataset was constructed using publicly available 10x Visium studies from each of the cancers identified using our atlas, namely BRCA^105^, BCC^106^, HCC^107^, NSCLC^108^, NPC^109^ and oral SCC (OSCC)^110^. The data was downloaded and processed to make it readable by the Seurat v5 R package^124^ for analysis. The SCTransform function in Seurat was used for normalisation and selection of variable features for each individual sample, returning all genes and 3000 variable features for downstream analyses. Integration was then conducted using the Harmony R package^125^. The final integrated dataset contained 122 samples from 69 individuals.

### Calculating and selecting CD4_Treg_8-high 10x Visium spots

To identify the spots which might contain CD4_Treg_8, the CD4_Treg_8 gene module was constructed using the tumour-specific Treg signature (*CXCL13, KLRB1, ADGRG1, BHLHE40, MYO7A, PTPN13, CSF1*) [Figure 2F] and some CD4 Treg-related genes (*CD4, IKZF4, FOXP3, IL2RA, CTLA4*). Seurat’s AddModuleScore function was then used to score the expression of the CD4_Treg_8 gene module in each spot. The top 1% of spots by score was selected for analysis [Figure S12B]. To ensure that the high module score was not skewed by a single (or a few) highly expressed genes, the coefficient of variation (CV) of all the genes in the CD4_Treg_8 module was calculated by taking the quotient of the standard deviation and the mean of the genes in the module. The top 1% of spots by score was then further filtered by taking the top 5% of spots with the most even distribution (lowest CV) of the module genes [Figure S12B], obtaining 1741 spots which we labelled as “CD4_Treg_8-high”. All the other spots were labelled as “NoTreg”.

### Annotating 10x Visium spots by microenvironment compartment with ESTIMATE

To determine the characteristics of the local microenvironment of each spot, the Estimation of STromal and Immune cells in MAlignant Tumours using Expression data (ESTIMATE) R package (version 1.0.13)^111^ was used. ESTIMATE infers the fraction of stromal and immune cells in each spot using gene expression signatures, and calculates a tumour purity score based on these fractions, enabling the classification of each spot into Immune_Hot, Stromal_Border or Tumour_Core regions.

To stratify all the spots according to these compartments together with CD4_Treg_8 status, spots were first separated according to their CD4_Treg_8 status (CD4_Treg_8-high vs NoTreg). This was to prevent the extreme population size imbalance (1741 CD4_Treg_8-high spots vs 219,620 NoTreg spots) from masking the classification of spots into each compartment. Within each respective group, each continuous ESTIMATE score was standardized into internally normalized Z-scores based on group-specific means and standard deviations to give a unified, variance-scaled distribution. Every spot was evaluated simultaneously across all three standardized metrics to assign its dominant local microenvironmental profile without sequential ordering bias, categorising each spot according to the metric displaying the highest Z-score. This generated six mutually exclusive microenvironmental compartments: Immune_Hot (highest relative immune cell infiltration), Stromal_Border (highest relative extracellular matrix or fibroblast presence), and Tumor_Core (highest relative malignant cell purity) for both CD4_Treg_8-infiltrated and CD4_Treg_8-absent niches.

---

**Supplementary Figure S1.**
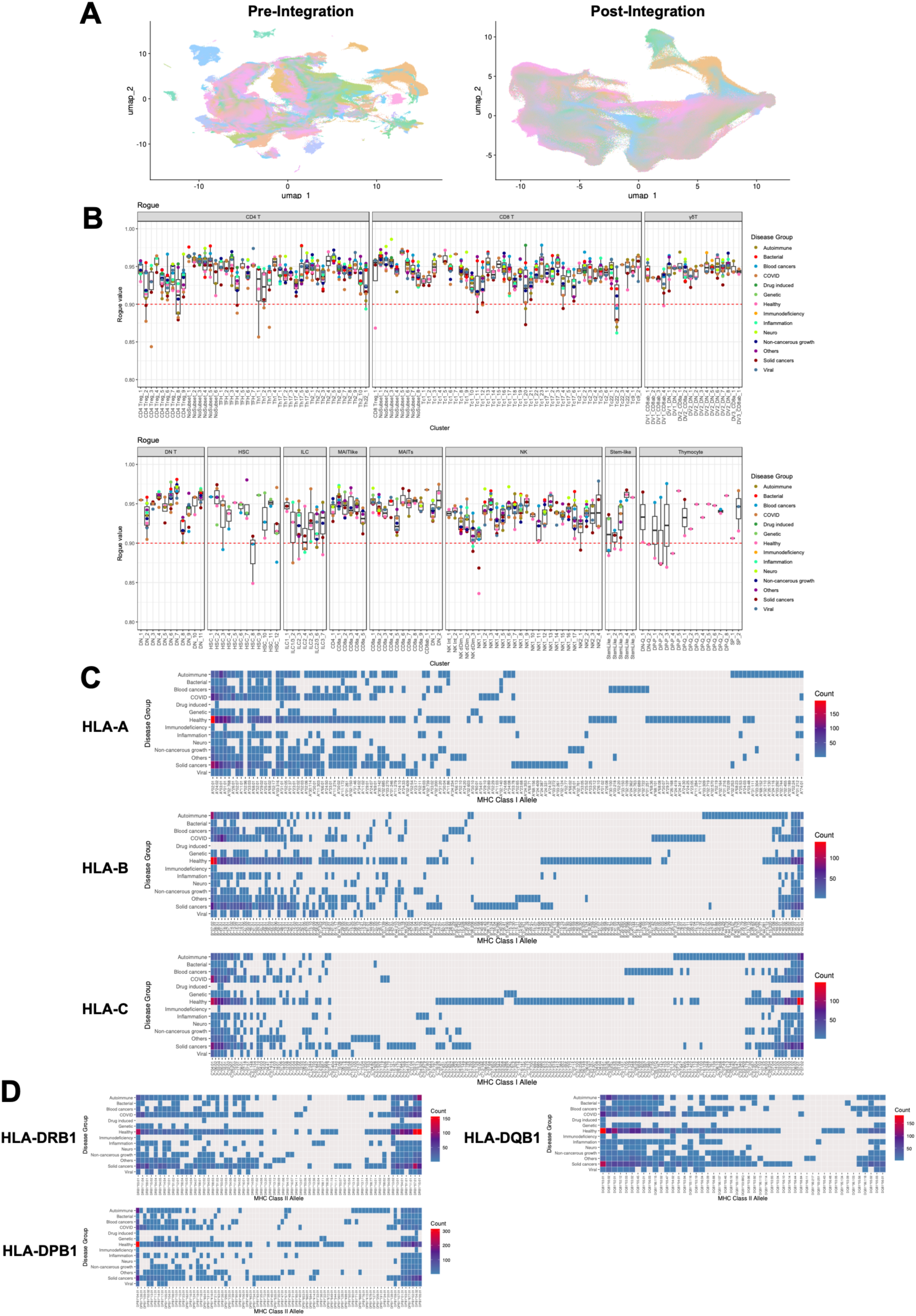
Construction of the T and ILC atlas (related to Figure 1). (**A**) UMAP visualisation of all cells in the atlas, pre- and post-integration, coloured by project. (**B**) Box plots showing the Rogue score of each cluster, grouping cells by disease group within each cluster to show homogeneity across disease contexts within each cluster. (**C**) Heatmap showing frequency of MHC Class I alleles within each disease group. (**D**) Heatmap showing frequency of MHC Class II alleles within each disease group.

**Supplementary Figure S2.**
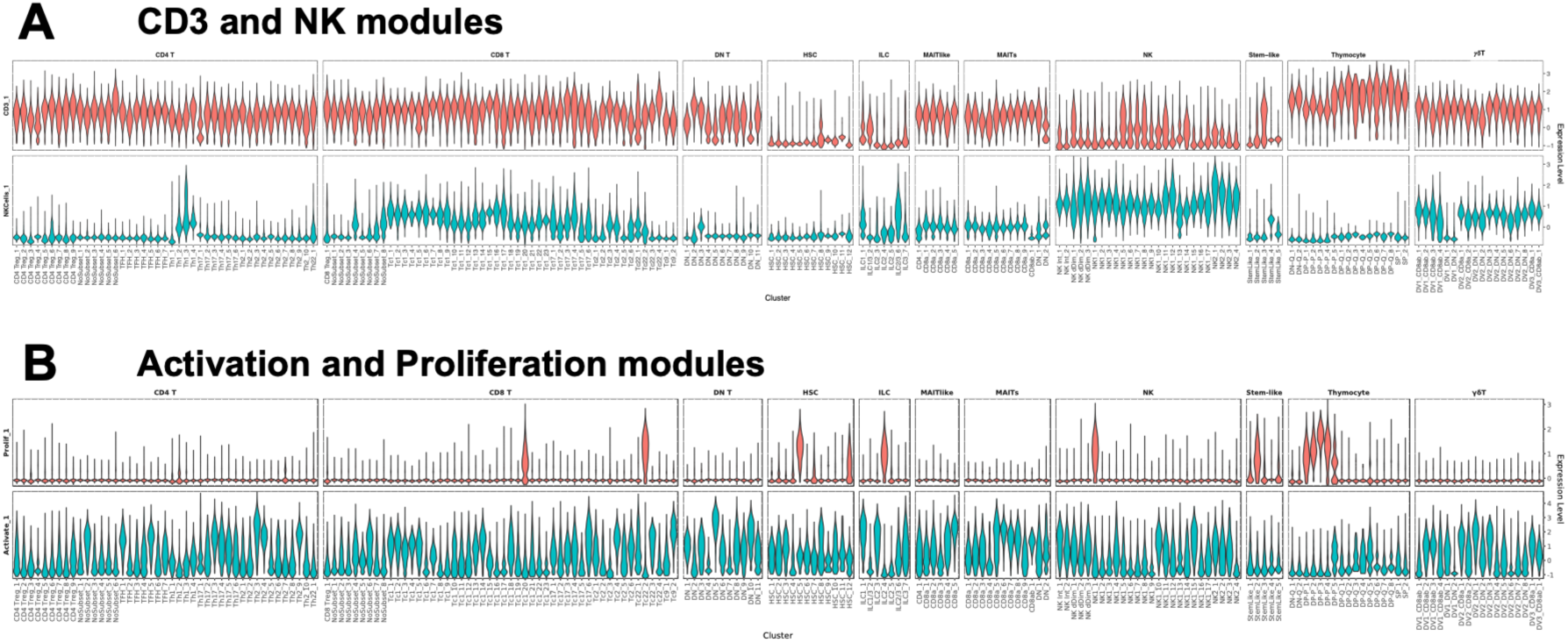
AddModuleScore analysis for general cluster annotation (related to Figure 1). (**A**) CD3 (*CD3E*, *CD3D* and *CD3G*) and NK (*KLRD1, KLRF1, NKG7, XCL1* and *XCL2*) modules. (**B**) Activation (*CD69, FOS* and *FOSB*) and the proliferation (*MKI67, TYMS* and *PCNA*) modules.

**Supplementary Figure S3.**
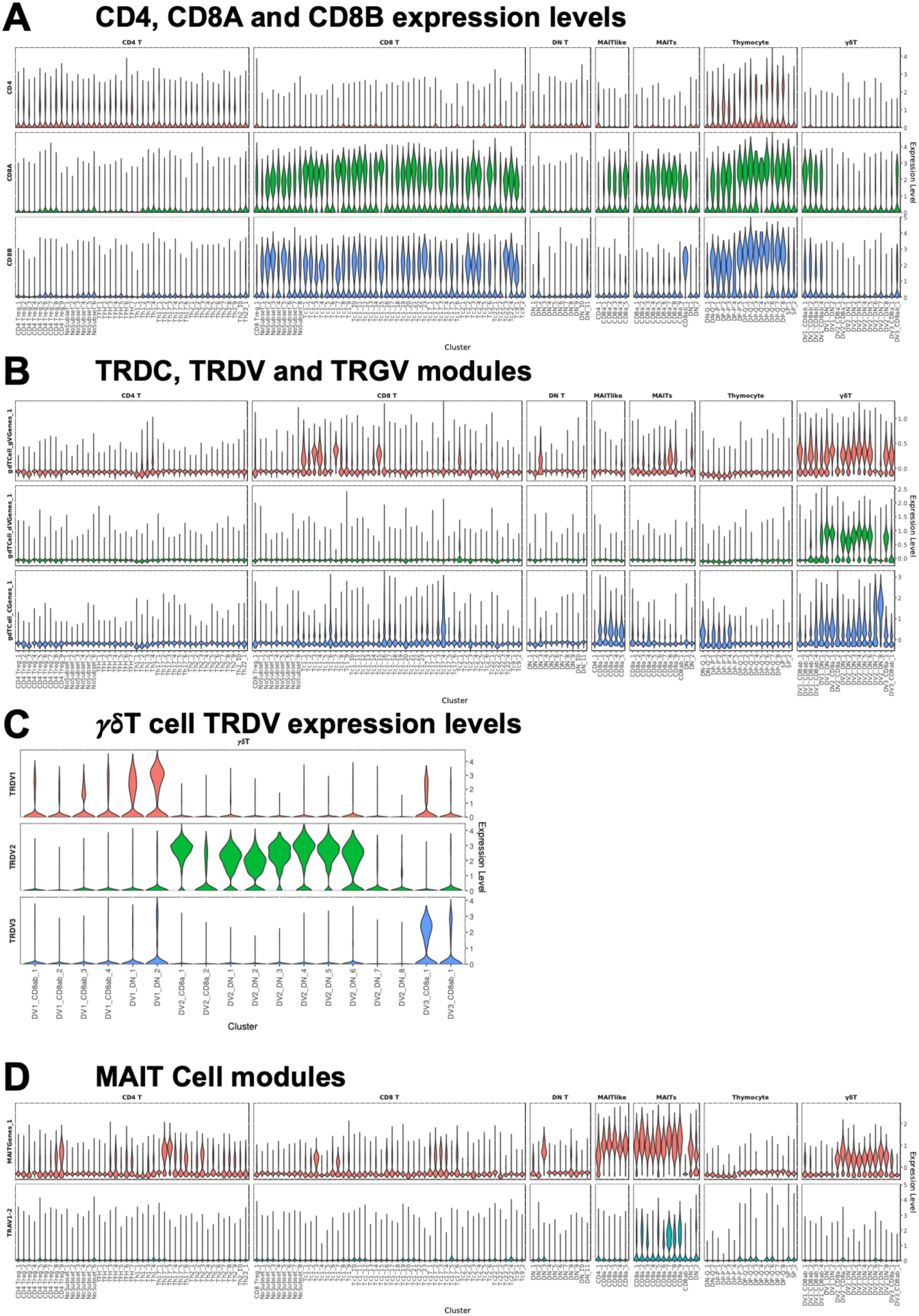
AddModuleScore analysis for T cell lineage annotation (related to Figures 2, 3 and 4). (**A**) *CD4*, *CD8A* and *CD8B* gene expression. (**B**) *γ*δT cell constant (*TRDC, TRGC1, TRGC2*), variable delta (*TRDV1, TRDV2, TRDV3*) and variable gamma (*TRGV2, TRGV3, TRGV4, TRGV5, TRGV8, TRGV9*) gene modules. (**C**) *γ*δT cell TRDV gene expression (**D**) MAIT cell (*TRAV1-2, KLRB1, SLC4A10*) gene modules.

**Supplementary Figure S4.**
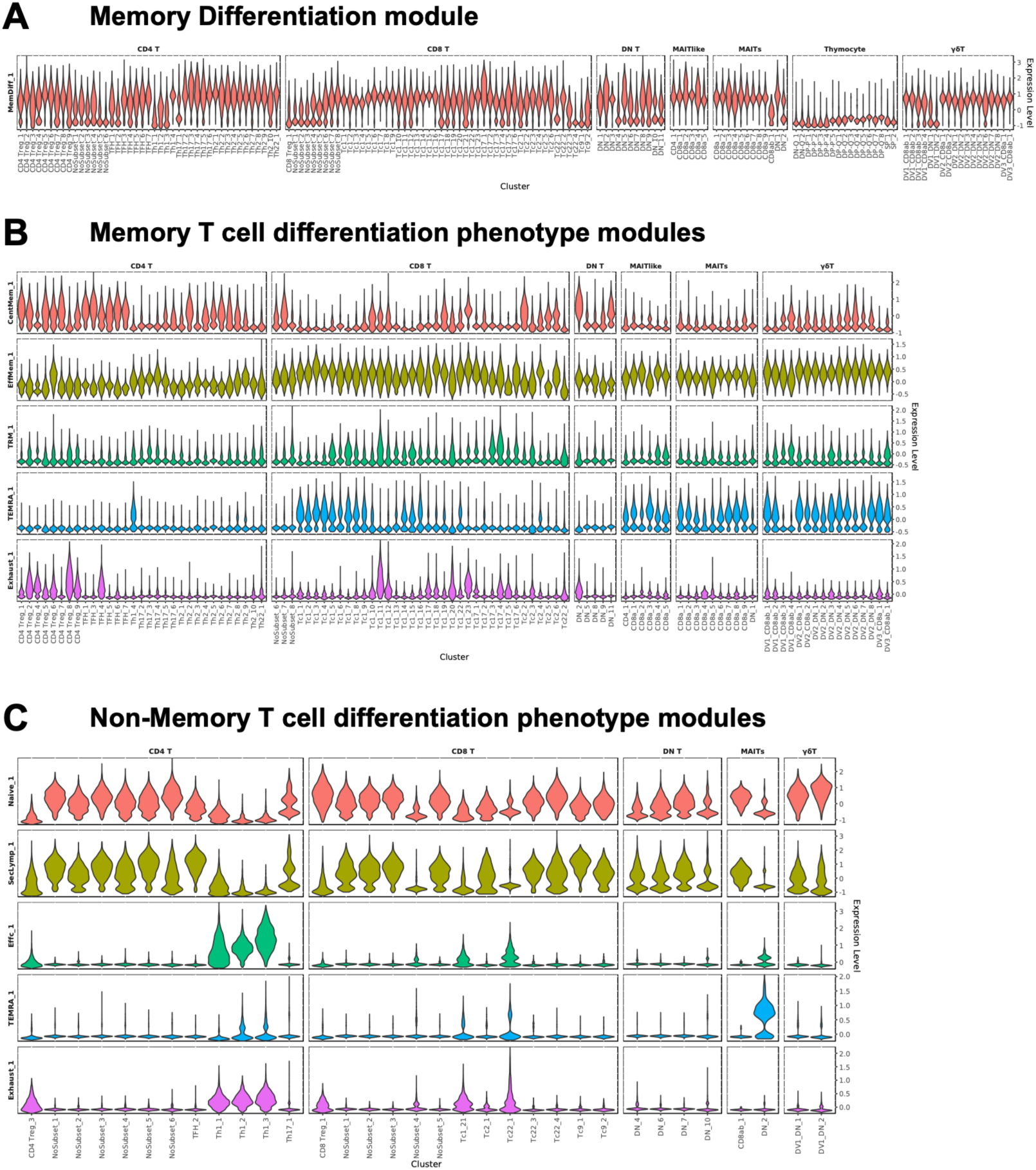
AddModuleScore analysis for T cell memory and differentiation annotation (related to Figures 2, 3 and 4). (**A**) Memory differentiation (*S100A4* and *GPR183*) module. (**B**) Differentiation modules for memory T cells, namely central memory (CM) (*CCR7, SELL* and *CD27*), effector memory (EM) (*TBX21, CX3CR1, CXCR3, CXCR4, FCGR3A, KLRD1* and *GZMK*), and tissue resident memory (TRM) (*ITGA1, ITGAE, PRDM1, ZNF683*) modules. (**C**) Differentiation modules for non-memory T cells, namely naïve (*TCF7, LEF1* and *CD27*), secondary lymphoid (*CCR7* and *SELL*) and effector (*CX3CR1, IFNG, PRF1, TNF, IL4* and *GZMB*) modules. Both (**B**) and (**C**) also have the terminal differentiation (TEMRA) (*KLRG1*, *S1PR5*, *B3GAT1*) and exhaustion (*CTLA4*, *PDCD1, TIGIT, ENTPD1, LAYN, HAVCR2, CXCL13*, *TNFRSF9* and *LAG3*) modules.

**Supplementary Figure S5.**
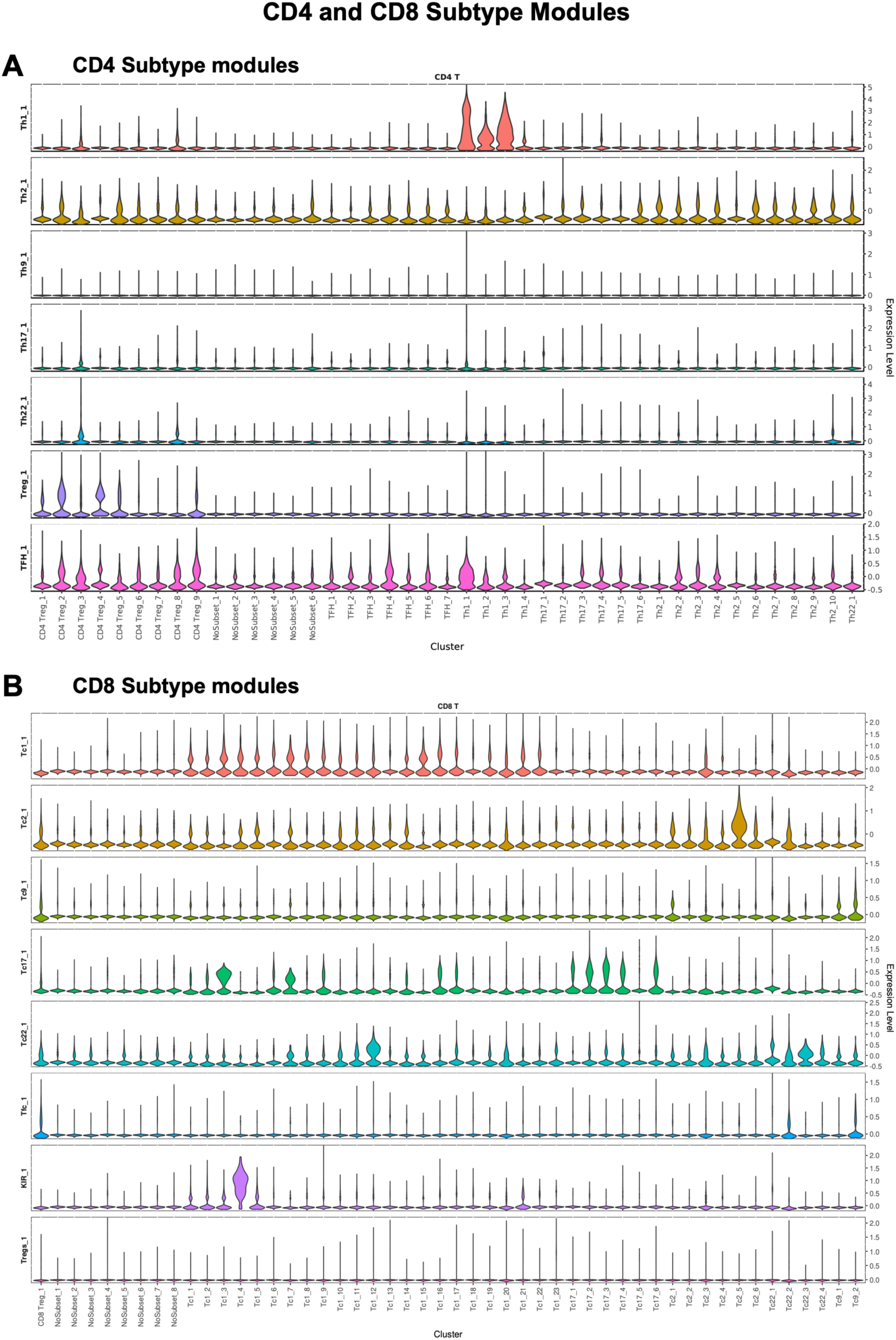
AddModuleScore analysis for CD4 and CD8 T cell subtype annotation (related to Figures 2 and 3). (**A**) CD4 subtype modules, namely T helper type (Th) 1 (*IFNG, TBX21*), Th2 (*GATA3, PTGDR2*), Th17 (*IL17A, RORC, RUNX1*), Th22 (*IL22, AHR*), T regulatory (Tregs) (*FOXP3, IL10*), T follicular helper (TFH) (*BCL6, CXCR5, ICOS*) and Th9 (*SPI1, IL9*). (**B**) CD8 subtype modules, namely Tc1 (*TBX21, EOMES*), Tc2 (*GATA3, PTGDR2*), Tc9 (*IRF4, STAT6, IL9*), Tc17 (*RORC, IL17A, KLRB1*), Tc22 (*Ahr, IL22, STAT1*), T follicular cytotoxic (Tfc) (*CXCR5, Bcl6, CD40LG*), killer cell immunoglobulin like receptor (KIR) expressing cells (*KIR3DL1, KIR2DL1, EOMES*) and CD8 Tregs (*FOXP3, IL10*).

**Supplementary Figure S6.**
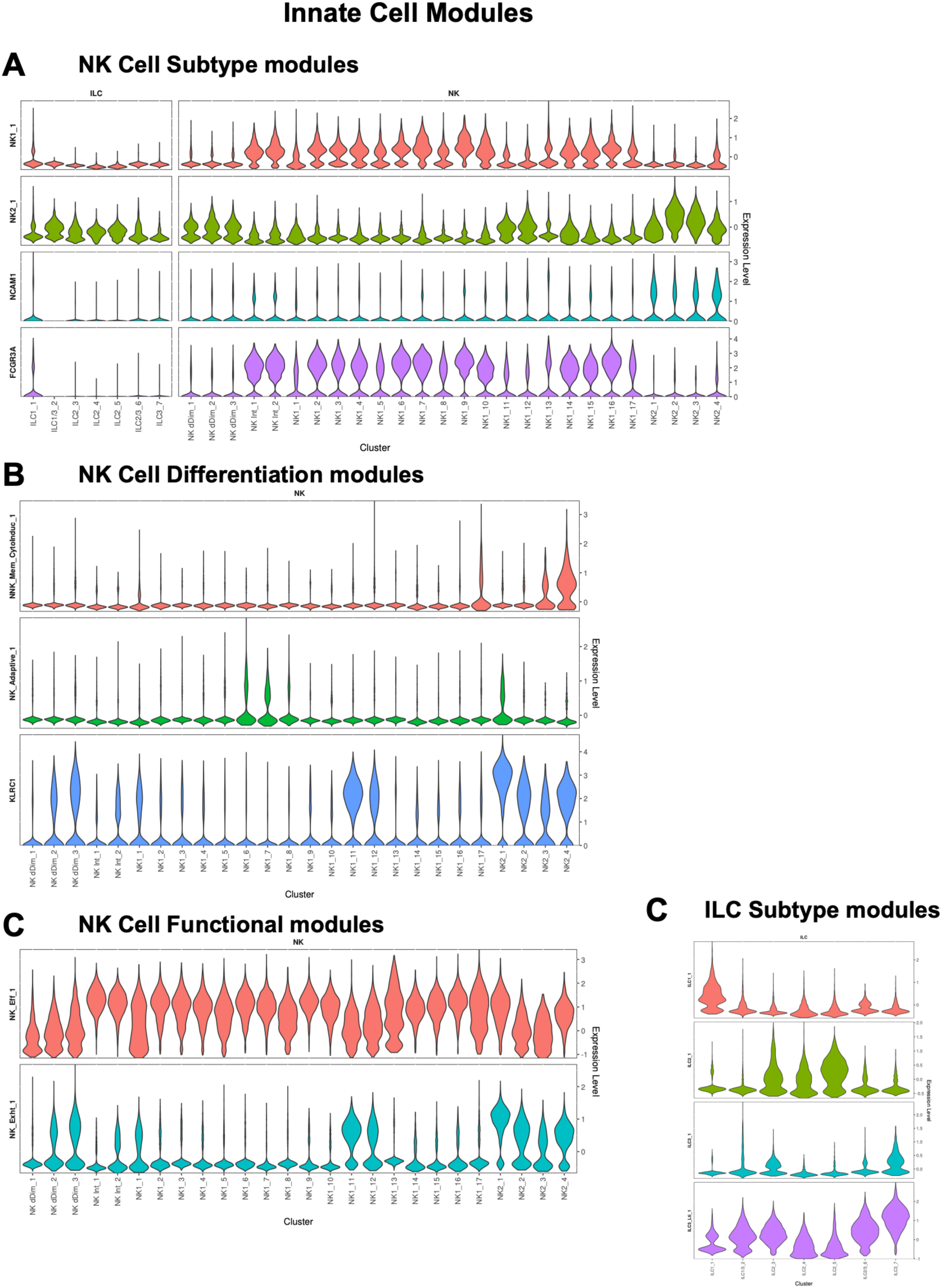
AddModuleScore analysis for NK and ILC subtype annotation (related to Figure 5). (**A**) NK cell subtype modules, namely NK1 (*FCGR3A, KIR3DL1, KIR3DL2*), NK2 (*NCAM1, SELL, IL2RA, IL18, GZMK*), and the genes *NCAM1* (CD56) and *FCGR3A* (CD16). (**B**) NK cell differentiation modules, namely memory-like cytokine induced NK cells (*TNFSF10* and *IL2RA*) and adaptive NK cells (*KLRC2* and *B3GAT1*), with the gene *KLRC1*. (**C**) NK cell functional modules, namely effector genes (*LAMP1, PRF1* and *GZMB*) and exhaustion or inhibitory markers (*PDCD1, KLRC1*). (**D**) ILC subset modules, namely ILC1 (*TBX21, IFNG, TNF, PRF1* and *IKZF3*), ILC2 (*GATA3, PTGDR2* and *IL1RL1*) and ILC3 (*RORC, AHR, IL23R* and *CCL20*). ILC3 were further annotated with a secondary subtype called the lymphoid tissue inducer (Lti) (*ID2, TNFSF11, LTB*).

**Supplementary Figure S7.**
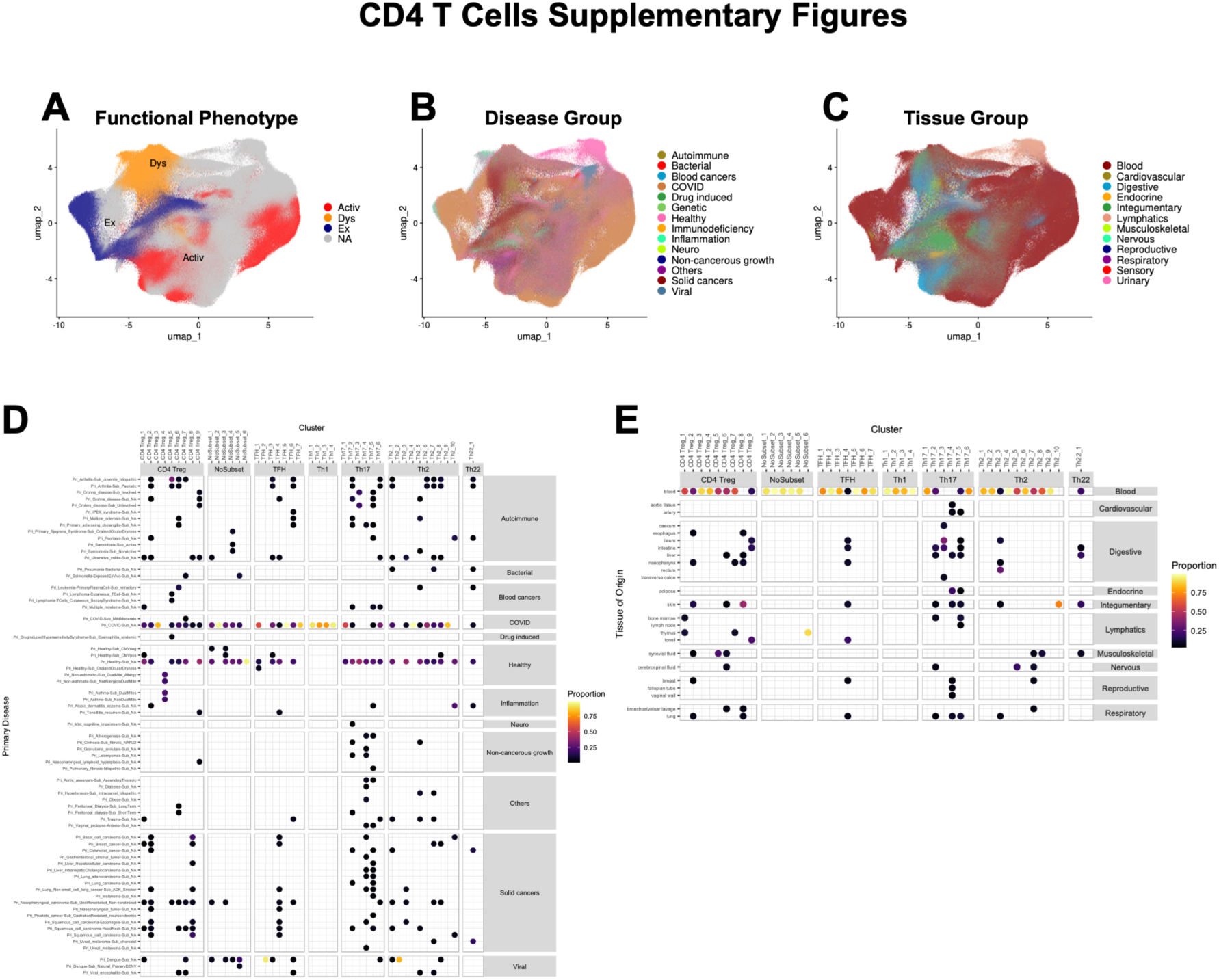
CD4 T cell compartment (related to Figure 2). (**A**) UMAP visualisation of CD4 clusters, coloured by functional phenotype. (**B**) UMAP visualisation of CD4 clusters, coloured by disease group. (**C**) UMAP visualisation of CD4 clusters, coloured by tissue group. (**D**) Dot plot heatmap showing the proportions of cells from the various primary disease subcategories within each cluster which add up to at least 70% of all the cells within the cluster. This shows the distribution of primary disease subcategories within each cluster. (**E**) Dot plot heatmap showing the proportions of cells from the various tissue origins within each cluster which add up to at least 70% of all the cells within the cluster. This shows the distribution of tissue origin within each cluster.

**Supplementary Figure S8.**
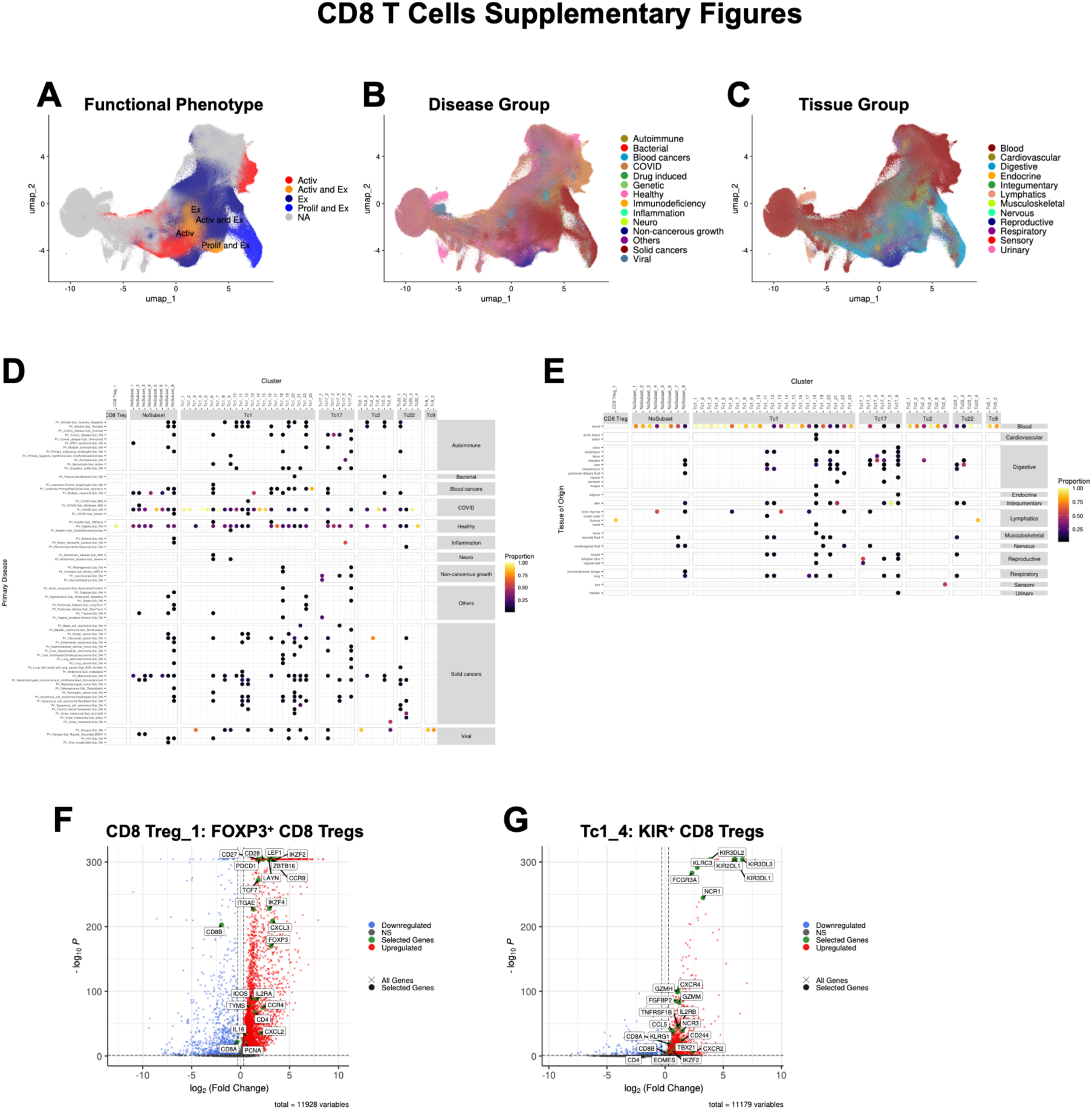
CD8 T cell compartment (related to Figure 3). (**A**) UMAP visualisation of CD8 clusters, coloured by functional phenotype. (**B**) UMAP visualisation of CD8 clusters, coloured by disease group. (**C**) UMAP visualisation of CD8 clusters, coloured by tissue group. (**D**) Dot plot heatmap showing the proportions of cells from the various primary disease subcategories within each cluster which add up to at least 70% of all the cells within the cluster. This shows the distribution of primary disease subcategories within each cluster. (**E**) Dot plot heatmap showing the proportions of cells from the various tissue origins within each cluster which add up to at least 70% of all the cells within the cluster. This shows the distribution of tissue origin within each cluster. (**F**) and (**G**) Volcano plots showing the DEGs of CD8 Treg clusters (CD8 Treg_1 and Tc1_4 respectively), calculated against CD8 T cells only.

**Supplementary Figure S9.**
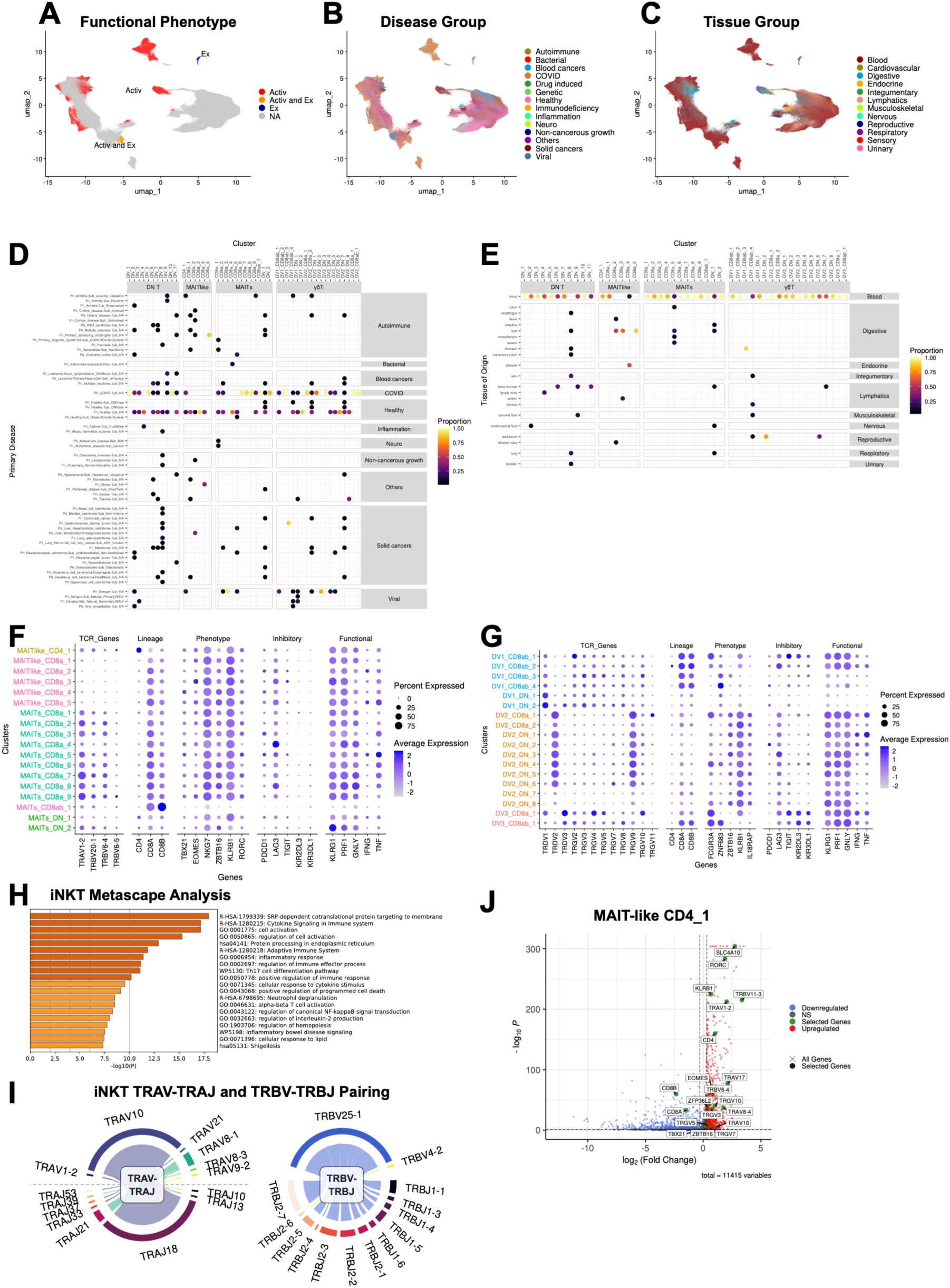
Unconventional T cell compartment (related to Figure 4). (**A**) UMAP visualisation of Unconventional T cell clusters, coloured by functional phenotype. (**B**) UMAP visualisation of Unconventional T cell clusters, coloured by disease group. (**C**) UMAP visualisation of Unconventional T cell clusters, coloured by tissue group. (**D**) Dot plot heatmap showing the proportions of cells from the various primary disease subcategories within each cluster which add up to at least 70% of all the cells within the cluster. This shows the distribution of primary disease subcategories within each cluster. (**E**) Dot plot heatmap showing the proportions of cells from the various tissue origins within each cluster which add up to at least 70% of all the cells within the cluster. This shows the distribution of tissue origin within each cluster. (**F**) Dot plot showing expression levels of selected genes for MAIT and MAIT-like cell clusters, grouped by co-receptor expression. (**G**) Dot plot showing expression levels of selected genes for *γ*δT cell clusters, grouped by subtype. (**H**) Top 20 enriched functional pathways in the iNKT cluster (DN_3) generated by Metascape, based on differentially-expressed genes (DEGs) calculated against all cells. (**I**) Circos plots showing paired TRAV-TRAJ and TRBV-TRBJ gene usage for the iNKT cluster (DN_3). (**J**) Volcano plot of the DEGs for the CD4^+^ MAIT-like cluster (MAIT-like CD4_1)

**Supplementary Figure S10.**
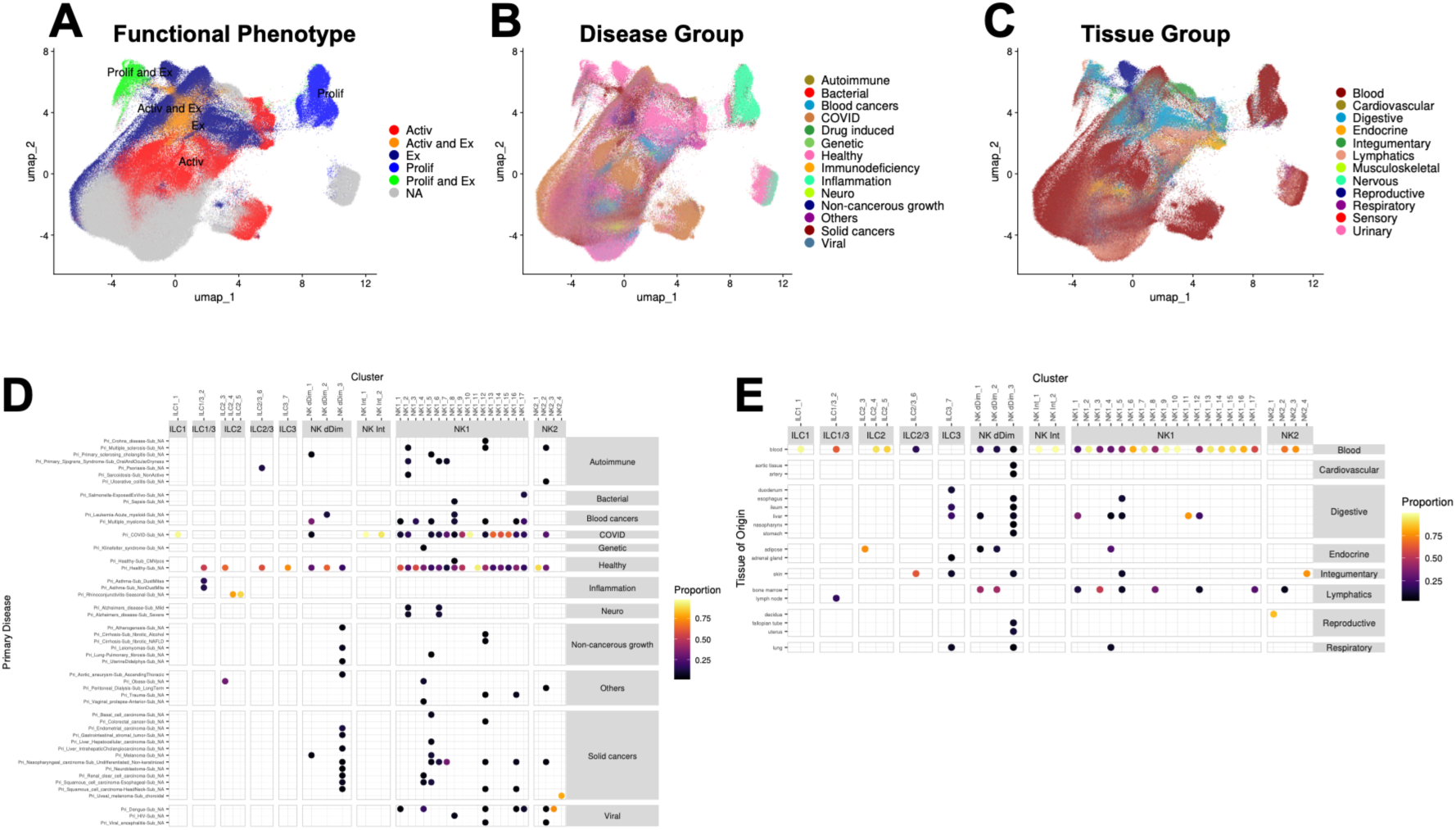
Innate cell compartment (related to Figure 5). (**A**) UMAP visualisation of Innate cell clusters, coloured by functional phenotype. (**B**) UMAP visualisation of Innate cell clusters, coloured by disease group. (**C**) UMAP visualisation of Innate cell clusters, coloured by tissue group. (**D**) Dot plot heatmap showing the proportions of cells from the various primary disease subcategories within each cluster which add up to at least 70% of all the cells within the cluster. This shows the distribution of primary disease subcategories within each cluster. (**E**) Dot plot heatmap showing the proportions of cells from the various tissue origins within each cluster which add up to at least 70% of all the cells within the cluster. This shows the distribution of tissue origin within each cluster.

**Supplementary Figure S11.**
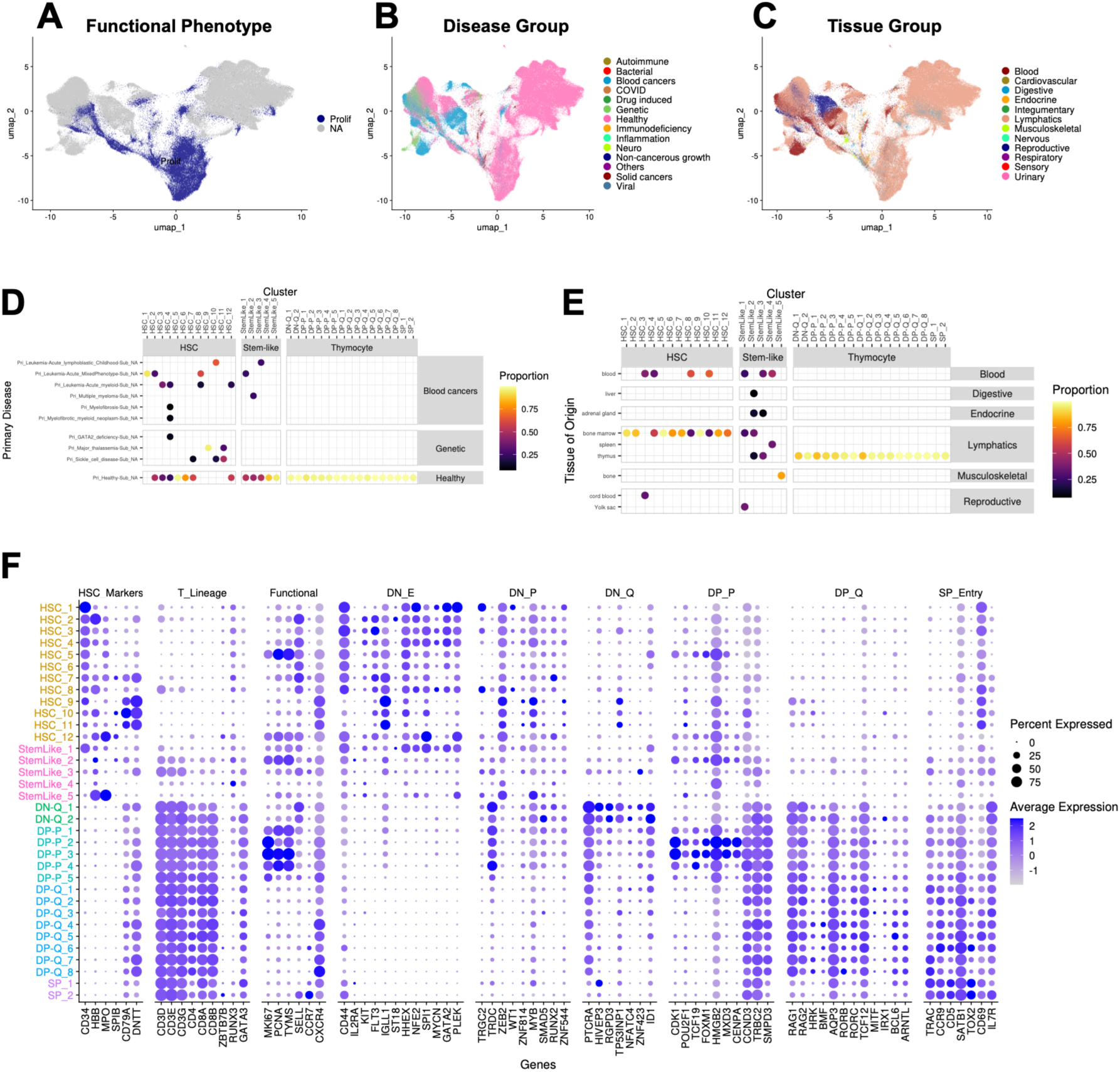
Immature cell compartment (related to Figure 6). (**A**) UMAP visualisation of Immature cell clusters, coloured by functional phenotype. (**B**) UMAP visualisation of Immature cell clusters, coloured by disease group. (**C**) UMAP visualisation of Immature cell clusters, coloured by tissue group. (**D**) Dot plot heatmap showing the proportions of cells from the various primary disease subcategories within each cluster which add up to at least 70% of all the cells within the cluster. This shows the distribution of primary disease subcategories within each cluster. (**E**) Dot plot heatmap showing the proportions of cells from the various tissue origins within each cluster which add up to at least 70% of all the cells within the cluster. This shows the distribution of tissue origin within each cluster. (**F**) Dot plot showing expression levels of selected genes for Immature cell clusters, grouped by main lineage and subtype.

**Supplementary Figure S12.**
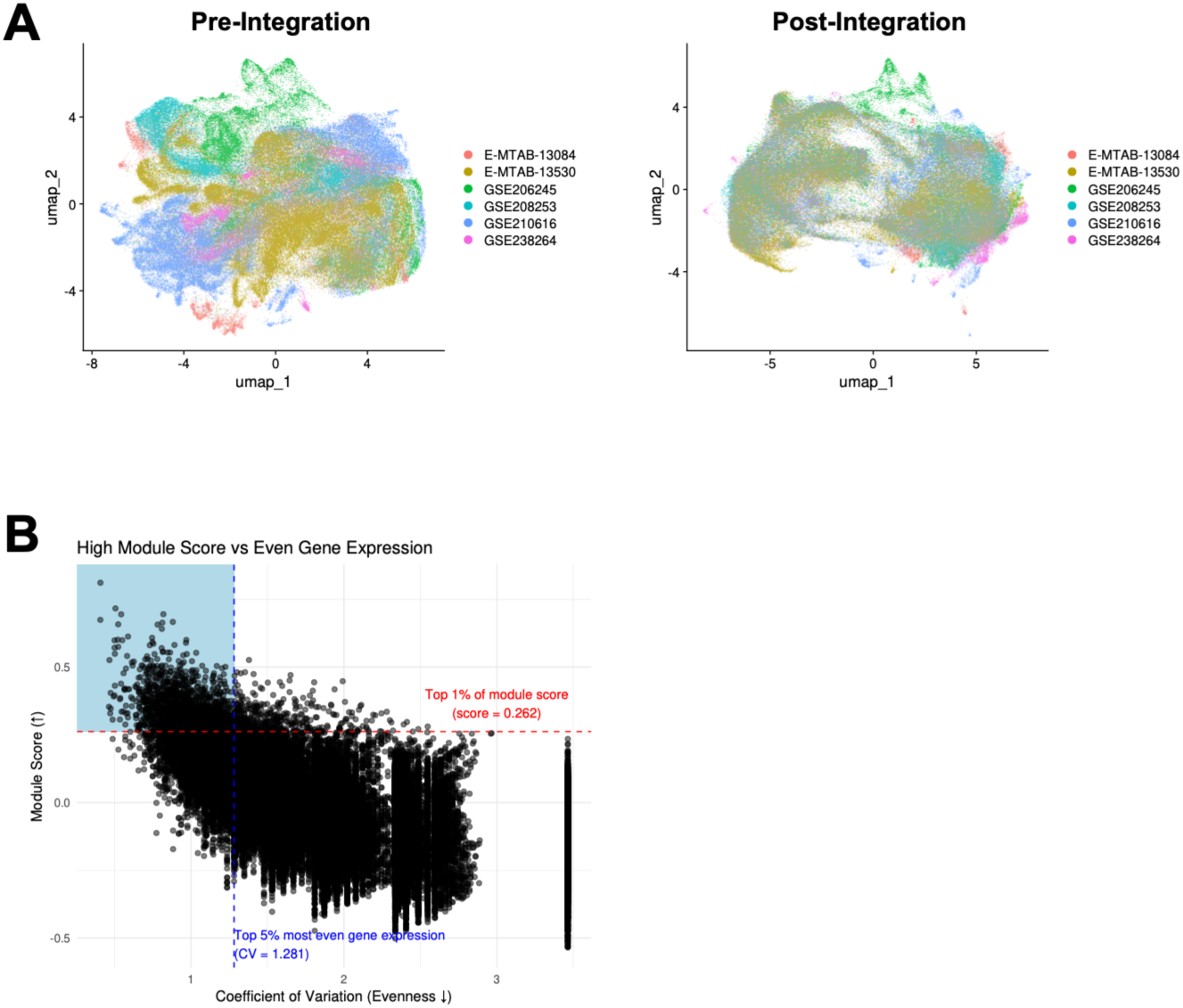
Analyses for spatial transcriptomics validation (related to Figure 7). (**A**) UMAP plots showing pre- and post-integration of the 10x Visium datasets, coloured by project. (**B**) Scatterplot describing the filtering criteria of spots to obtain CD4_Treg_8-high spots. The red dotted line shows the top 1% of spots by CD4_Treg_8 gene module score, while the blue dotted line shows the top 5% of spots by the most even gene expression as determined by coefficient of variation.

